# *Amblyomma mixtum* free-living stages: Summer and winter use, preference, and niche width in an agroecosystem (Yopal, Casanare, Colombia)

**DOI:** 10.1101/2020.12.23.424125

**Authors:** Elkin Forero-Becerra, Alberto Acosta, Efraín Benavides, Marylin Hidalgo

## Abstract

Studying a species’ tolerance to an ecosystem’s environmental conditions and its selection of available resources is relevant in ecological and evolutionary terms. Moreover, formulation of effective control strategies implicitly includes the study of habitat use and preference and niche width in anthropogenically transformed natural landscapes. Here, we evaluated whether the use, habitat preference, and niche range of the *Amblyomma mixtum* tick changed between stages, habitats, and seasons (summer-winter 2019) on a farm in Yopal (Casanare, Colombia). To this end, the presence and relative abundance of larvae, nymphs, and free-living adults was quantified in four different habitats according to the type of vegetation cover (Riparian Forest, Cocoa Crop, King Grass Crop, and Star Grass Paddock). Habitat availability was calculated, environmental variables were analyzed, and various indices of habitat use and preference and niche width were calculated. *A. mixtum*’s habitat use and preference and niche width changed between stages, habitat types, and time of the year. The total abundance of *A. mixtum* was an order of magnitude greater in summer than winter. Nymphs and larvae dominated it in the summer and adults in the winter. In summer, all the stages used the four habitats. In winter, the larvae did not use two habitats (Riparian Forest and Cocoa Crop); nymphs did not use the cocoa crop. *A. mixtum* adults used all the habitats in both seasons. In summer, the nymphs and larvae preferred three of the four habitats (King Grass Crop, Star Grass Paddock, and Cocoa Crops), while adults preferred the King Grass Crop. In winter, the nymphs and larvae preferred the King Grass Crop and Star Grass Paddock, while the adults preferred the King Grass Crop. The value of the niche width index was high for larvae, nymphs, and adults in summer, while it was high only for adults in winter. *A. mixtum* is exposed to significant daily, seasonal, and multiannual variations in relative humidity (minimum 30%), ambient temperature (minimum 18°C), solar radiation (maximum 800 W/m^2^), and precipitation (maximum 481 mm/month). Thus, the local *A. mixtum* population could rapidly acclimatize to changing habitats (unstable or temporary) under fluctuating environmental conditions (e.g., King Grass Crop). However, the winter flood season in Yopal could exceed *A. mixtum*’s adaptive capacity during its most vulnerable stages. Mathematically, a low number of female *A. mixtum*, surviving the most demanding environmental conditions, could sufficiently ensure the population’s persistence, which, coupled with the vast host range, could facilitate the ticks stages’ dispersal among habitats to complete their life cycle. *A. mixtum*’s population control should be carried out during its season of greater vulnerability (winter), when the population is low, particularly the females.

## Introduction

Animals’ habitat use, preference, niche width, and resource selection allows researchers to explain abundance, spatial distribution (occurrence maps for conservation or health planning purposes), and evolutionary processes (cost and benefits of differential habitat use, adaptations, niche separation, and speciation) of a given organism-species to infer its ecological requirements (animal–habitat relationships) in a changing world, including the effects of human activities (habitat loss) on wildlife’s ability to colonize and use specific areas [1–3].

Habitat is defined as the place with all the conditions and resources necessary for survival, development, reproduction, and the establishment of local populations [4]. Habitat is also the spatially limited site where abundance or other population parameters (rate of growth = R) differ from those of other localities or contiguous patches, or habitats [5]. For example, in host-parasite relationships, as in ticks, the host is both the space and habitat that offers the resource (e.g., blood to the parasite). Different host types will define the tick’s population abundance. However, this does not mean that all the available hosts in the habitat will be used. Resources are defined by [6] as a discrete unit (attributes or environmental variables), such as forest, grassland, or crop categories, within the vegetation cover types that an animal may encounter, use, and discriminate. Thus, an animal will instinctively determine, based on cost-benefit, which category of any particular resource should be used. A used resource unit is the one that has received some investment by an animal, including time-energy expended, distance traveled, or residence time in the unit-habitat [3].

However, the resource, as part of the organism’s habitat, can be available or unavailable. An available resource unit, or one spatially limited by the researcher, is one that can potentially be encountered by an animal [3], depending on the presence or absence of limiting factors, such as physical and biological (parasitism, predation, and competition) in time and space, that could prevent the establishment, survival, development, or reproduction of any given organism [2]. If such an animal is not found, the target resource and the habitat is considered unused. Availability is similarly defined by the proportional occurrence (area) of discrete levels of a resource; that is, the area covered by different vegetation types or habitats. In resource selection studies, it is important to define availability because it determines the spatial distribution of resources in those habitats against which use is compared [7]. Comparing used to unused resource units, or other used units, to available resource units allows inferences regarding selection and preference [3]. Habitat preference, on the other hand, is understood as a consequence of habitat selection by a reproductive female, larvae, nymph or adult, or the non-random asymmetric use of any resource(s) by each individual stage in the population, over other possible resources found in other available habitats [5]. Habitat preference must be inferred at the population level. It results from many individuals choosing the same physical or biological resource in a habitat (i.e. refuge), even over the availability of others [2].

However, the asymmetric use of any resource(s) in the habitat depends on the organism’s requirements. It also varies in time and space in relation to life cycle stages or a change in resources [3,8]. Also, it depends on the quality, quantity, and availability of resources in a particular habitat [9]. Essentially, an animal’s habitat preference is assessed by its relative population abundance in all the habitats it uses, including their resources, according to the availability of comparable habitats (area of potential use with resources an animal can encounter). As stated by some authors [2,10], this implies establishing realistic biological categories depending on the organisms’ life history (larvae, nymphs, adults) and the discrete habitats (resources) that the organisms may use (grass, crops, forest), not use, select, and prefer (asymmetric use) at the population level.

On the other hand, the concept of niche width refers essentially to the diversity of resources used and conditions needed in a habitat by any organism or group of organisms [11]. In other words, the way a species exists in the habitat using resources (hiding, hunting, moving, and parasitizing) and the habitat conditions (environmental factors, like site quality, temperature, and humidity) in which organisms occur and how such conditions affect it (survival, diapause, death). Thus, species abundance and spatial distribution are attributed to its niche amplitude (both as a proxy) [11]. A niche is one specie’s multidimensional space, where a measurement of its dimensions (one or the set of environmental factors) is called niche width, breath, or amplitude [12]. In any habitat, each factor can have a lower or upper limit. Anything below or above those limits will hinder the species from growing or cause its elimination (exceeding its tolerance limits).

These limits are the optimum range for species performance, survival, and positive population growth rate. [13] indicated that a factor’s range between minimum and maximum values represents the specie’s ecological amplitude for that factor. Species with wide niche breadth are highly adaptable. They can use a broader spectrum of resources, even with limited efficiency, according to scarcity or availability of those resources in the habitat. They can adjust easily (through morphological, physiological, and behavioral traits) to changes in the environment. These species could potentially live in a broad range of temperature or humidity conditions, and they can also attain higher population abundances and occupy several different habitats (resource possibilities).

These species are referred to as generalist species and include several *Amblyomma* tick species. Specialist species have a narrow niche breadth [14]. Habitat quality determines a species’ performance (presence, abundance, fitness, and growth rate) related to its niche breadth. In turn, the species’ performance (high or low abundance) could indicate the habitat’s attributes (good or bad [13]). Although it is difficult to describe and find any specie’s ecological niche (infinite niche dimensions), few significant variables and the species response to each of them (survival, abundance) may be sufficient to measure its established niche and infer its niche width [11].

Therefore, a specie’s selection of one or more habitats throughout its life cycle (stages) is essential to the population’s preservation and persistence. Biotic and abiotic factors and habitat resources affect survival, reproduction, population growth rate, and spatial distribution at different time and space scales, evolutionary and/or ecological. Depending on the species’ niche, individuals actively seek out and select habitats that provide an adequate range of conditions and resources [15]. However, natural habitats are constantly changing, and so are the species. Through evolutionary processes, they can adapt their morphology, physiology, and behavior to use new habitats, even managing to expand their fundamental ecological niche to unstable and dynamic habitats [16].

Human activities, for instance, benefit synanthropic species’ spatial distribution and abundance [17] by expanding their established niche [18], particularly in agroecosystems [19]. The favored species (like some ticks) increase their dispersion capacity, colonizing different habitats, and decreasing their extinction probability [20]. Forest degradation and fragmentation, for example, have prompted the occurrence of *Amblyomma aureolatum* in urban areas of the city of São Paulo (Brazil) [21,22].

Several field studies have evaluated *Amblyomma* ticks’ seasonal population dynamics and abundance (see [23–29]). However, few have approached the use of resources in the habitat or the inference of their niche range on a local scale. That is, *A. mixtum*’s preference for various habitats in a single agroecosystem, the role that the domestic and wild host community would play there, the competition for resources from those habitats with other ticks, and the associated dynamics of rickettsial infection in local animals and humans are not known. Some examples of research on ixodids of vector-pathogen-host relationships are *Ixodes scapularis* [30]; *Amblyomma fuscum* [31]; *Amblyomma americanum* [32]; *Amblyomma aureolatum* and *Amblyomma ovale* [33]; *Amblyomma triste* [34]; *Ixodes ricinus* [35]; *Ixodes pacificus* [36]; *Amblyomma tuberculatum* [37]; *Rhipicephalus microplus* [38], and *Amblyomma maculatum* [39].

Although some factors that could affect *Amblyomma mixtum*’s habitat use have been studied at the local [40] or laboratory level [41], or modeled on a regional [42] or continental scale [43], there is still a lack of empirical information that evaluates and integrates aspects of this tick’s use, habitat preference, and niche range in an agricultural system and whether the non-parasitic stages and population respond differently over time (seasonality). Likewise, there are no studies in Colombia that clearly indicate *A. mixtum*’s limit of altitudinal distribution in the Eastern Cordillera or explanations based on empirical data of its absence in the Bogota savannah. There is a trend towards studying the tick’s eco-epidemiology, particularly the limits of their geographic distribution, using mathematical modelings of their niche based on records of local presence and seasonal abundance [44]. However, the absence of observations cannot be interpreted as the specie’s non-existence in the site [45].

These models are used to obtain mechanistic explanations of vectors and pathogens’ dynamics or predict transmission risks in space and time. However, detailed empirical information is needed to add biological realism levels to understand that not all ticks and their hosts behave the same way in all environments [46]. Therefore, obtaining empirical information on a tick species’ life history, its tolerance to extreme abiotic variables values, and its interaction with abiotic factors is crucial for decision making about the management of the vector and its impact on public health.

Ticks of the *Amblyomma* genus have been recognized as vectors of rickettsial agents in a number of countries; thus, the importance of their study [47]. In Colombia, some species, particularly those included in the *Amblyomma cajennense* species complex (see [48]), have been reported infected with *Rickettsia rickettsii*. Such is the case of *Amblyomma patinoi* (Villeta, Cundinamarca [49]), *A. cajennense* sensu lato (La Sierra and Rosas, Cauca, [50]), and *Amblyomma mixtum* in several regions, including Yopal (Casanare [51]).

According to a recent epidemiological study in the Matepantano farm in Yopal, Casanare (Hidalgo, et al., unpublished information), there is a high proportion of the human population (92%, 77/84) with antibody titers to *Rickettsia* spp. due to their exposure to ticks. Considering the scarce information existing between non-parasitic *A. mixtum’s* stages and its relationship with resources and habitat conditions, we investigated the population’s use and preference of four different habitats and their niche width. We compared the summer (February) and winter (August) to see if, as had been suggested [3], the seasonal variation (precipitation, humidity, and temperature) could altered the use and preference of the habitats and the niche width (intraspecific variation).

## Materials and methods

### Habitat selection and sampling details

The work was carried out within what we have called Matepantano farm, an area surrounding part of the main installations of the Hacienda Matepantano (Fig 1), a private-owned land where academic agricultural spaces and activities occur (Vereda Matepantano, Yopal, Casanare, Colombia). Non-protected species were sampled. The field collections were possible thanks to the permissions of the Ministry of the Environment of Colombia (001-2018). The farm is located at an altitude between 256-270 m.a.s.l. It is delimited by gallery forest corridors, which are home to wildlife. Initially, hundreds of successive aerial images, taken with a digital camera mounted on a Phantom 4 Advanced DJI drone from an altitude of 150 m (S1 Appendix), were used to create an orthomosaic of the study area (domain) in ArcGIS Desktop^®^ (Esri, Redlands, CA, USA).

**Fig. 1.**
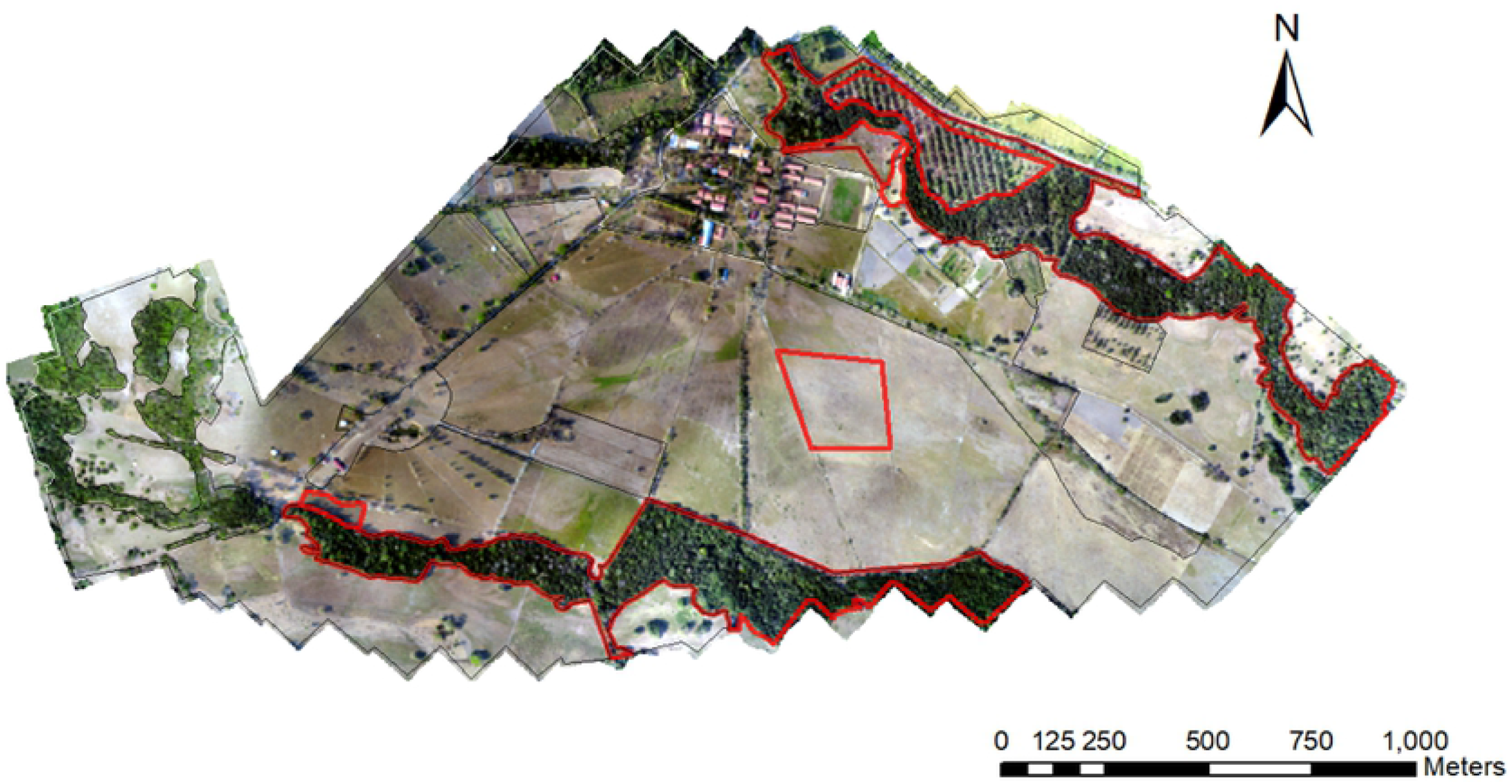
Aerial view of the study area at the Matepantano farm. Red lines encircled areas of the selected habitats.

The spatial scale was determined to allow us to capture the domain’s maximum spatial heterogeneity (*discrete vegetation types*) caused by anthropic intervention processes, like livestock, crops, and pastures, to see their effect on *A. mixtum*’s use of resources. The different habitats were defined and delimited on the orthomosaic according to the type of vegetation and the different patches of vegetation within each habitat (Fig 2). Then, ArcGIS was used to calculate the total area of the Matepantano farm (3’203.786.9 m^2^) and the area of each of the four selected habitats. Only those areas (discrete patches or paddocks) of habitat where tick sampling was conducted were considered (previously confirmed presence of *A. mixtum* in pre-samples).

**Fig. 2.**
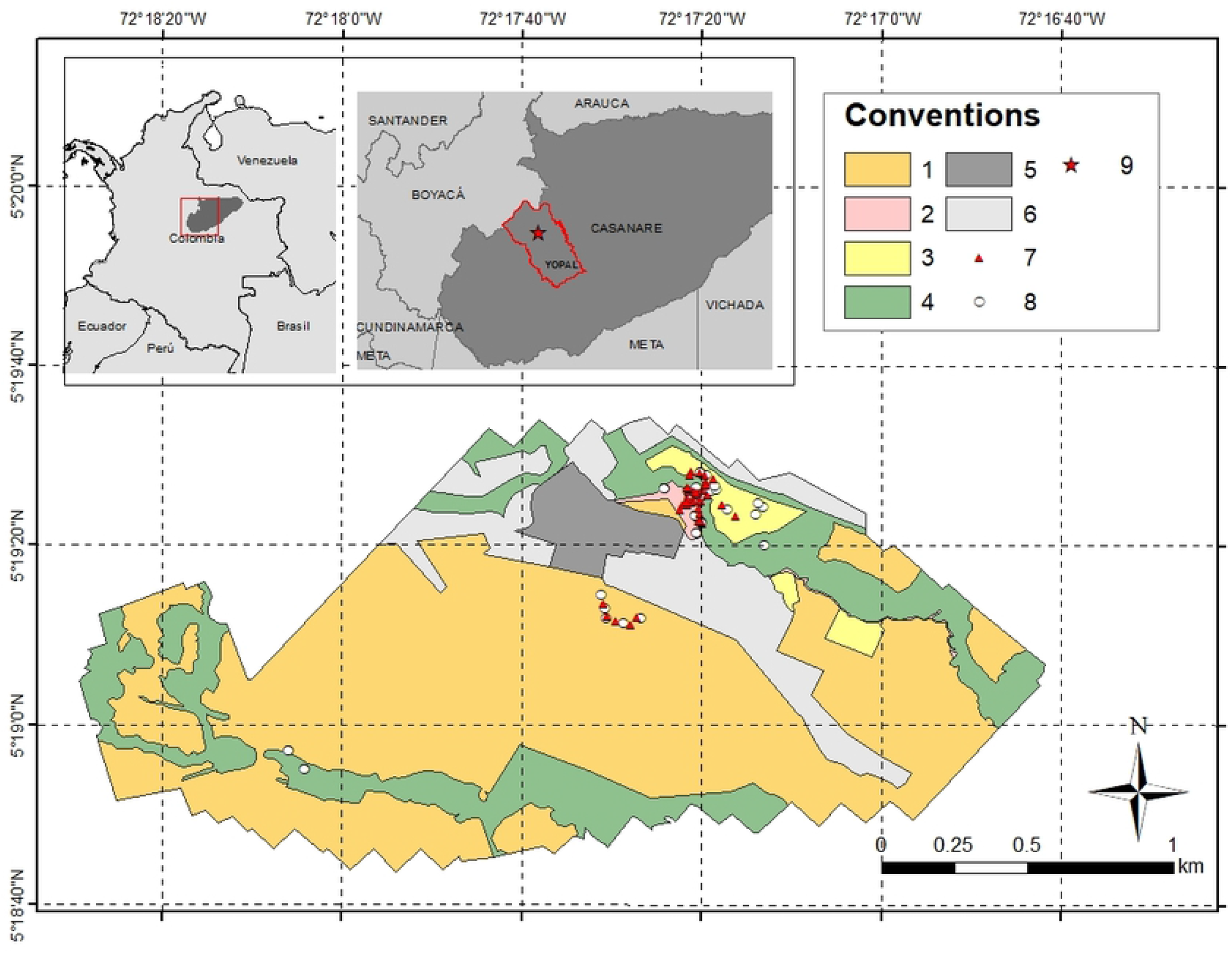
Map of the study area at the Matepantano farm and the four selected habitats. The study area is about 3,2 km^2^ representing a 6% of the total area of Matepantano estate. Conventions: 1-Star Grass paddocks (1’955,090 m^2^); 2-King Grass crop (22,416 m^2^); 3-Cocoa crop (81,341 m^2^); 4-Riparian forest (646,099 m^2^); 5-Main buildings area (112,833 m^2^); 6-Other area (386,007 m^2^); 7-CO_2_ traps; 8-Dragging by cloth transects; 9-Regional localization of the Matepantano farm. Note: All CO_2_ traps and dragging by cloths transects with GPS data for both seasons, summer and winter, are represented.

The areas finally selected were the following: riparian forest, 450.552 m^2^ of a total of 646.099.0 m^2^; cocoa crop (*Theobroma cacao*), 59.940 m^2^ of a total of 81.341.3 m^2^; Star Grass paddock, 54.968 m^2^ of a total of 1’955.090.4 m^2^; and King Grass crop, 13.245 m^2^ of a total of 22.416.3 m^2^. For this study, these four areas were called “available area”; that is, areas with potential to be used by the *A. mixtum* tick population. This available area is essential in determining the observed use in each habitat (presence and abundance of ticks) and defining habitat preference and niche width (indices applied). All four habitats were equally represented in the habitat availability area, except for the ‘Star Grass paddock,’ which we did not want to over-represent in the habitat availability, given its higher proportion in the domain area (Fig 2).

The mathematical calculations (indices) used the same values for summer and winter in the four available areas, given that the same points were sampled in time. Within the Matepantano farm, some areas were not sampled, such as buildings for academic, recreational, food, and residential activities and some discrete patches of the four habitats (386.007.0 m^2^) (Fig 2). A sampling of free-living ticks within each habitat was performed from February 8^th^ to 12^th^, 2019, during the summer season (November-March) and from August 16^th^ to 19^th^, 2019, during the winter season (April-October). Seasonality alters the relationship between the tick and the use of the resource-habitat [3]. Within each habitat category, the CO_2_ trap and white flannel dragging transect sampling units (grouping all the individuals captured by trap or transect) were located at convenience, following biological criteria (biology and pattern of tick activity and the probability of being a human vector), with replicates for each habitat (where abundance and presence-use response variables were measured). The habitat fragments, patches, and paddocks, chosen and sampled from the photomosaic, favored those within a radius of approximately 500 m from the buildings of the Matepantano farm, including the places frequently used for work and agropastoral practices (Figs 1 and 2). The number of traps and transects were fairly similar between habitats to facilitate comparison of relative abundances. Random sampling was not used for the location of the sampling units (trap or transect) in the vegetation types because, as indicated by other authors [3], this usually results in few or no observations of habitat use.

### Generalities of the compared habitats

#### **Riparian Forest** (5.32224° N and 72.28696° W)

This habitat was a gallery forest bordered by a permanent creek and dominated by adults and juveniles of the *Attalea butyracea* (Arecaceae) palm, a closed canopy, and an open understory (Table 1). There was no apparent change in vegetation structure between summer and winter (stable structure), with saturated soil in winter and litter in both seasons. Tick abundance sampling was conducted between 2 and 8 m into the forest, outside the edge zone.

**Table 1.** Structural characteristics of the four habitat selected where ticks were collected.

|  | Riparian forest | Cocoa crop | King Grass crop | Star Grass paddocks |
| --- | --- | --- | --- | --- |
| **Dominant terrestrial vegetation** | Palm<br>(*Attalea butyracea*).<br>Poor litter with presence of nuts, tree branches, litter debris and fungi. | Cocoa, acacia, banana trees. Acacia leaves dominates the litter with fruits and tree branches. | King grass dominates this area with some scattered trees and thorny bushes. There is no litter. | Star grass (*Cynodon* sp.) and kermes oak. Litter was scarce with predominance of grass dry leaves. Soil is floodable by rains, but humidity was found during a 24 h period. |
| **Natural light availability (%)** | 25 | 50 | 100 | 100 |
| **Number of vegetation strata** | 4 | 3 | 1-2 | 2 |
| **Litter depth (cm)** | 2,9 ($\pm 1,6$) | 6,8 ($\pm 3,5$) | 8,5 ($\pm 2,2$) | 2,8 ($\pm 2,8$) |
| **Litter humidity (%) in summer** | 56,4 ($\pm 6,5$) | 47,4 ($\pm 5,1$) | 62,9 ($\pm 8,2$) | 39,9 ($\pm 6,7$) |
| **Soil dominant fraction** | Clay soil with some sand. | Clay soil (alfisol type).<br>Arcilloso – tipo alfisol. There are rocks at deeper digging.<br>The soil has acidic pH. | Clay soil with few silt and fine sand soils. | Clay soil. |

#### **Cocoa Crop** (5.32312° N and 72.28691° W)

This habitat was a cacao crop (*Theobroma cacao*). The farm used the *tresbolillo* technique to plant the cocoa. This technique entails locating the plants in a triangular shape with a distance of 3.5 x 3.5 m between them. The crop was mixed with *Acacia* trees to provide shade (Table 1), in addition to some banana plants. In summer, the crop soil was dry (cracked), with leaf litter and dehydrated plants caused by a couple of months of drought (december-january). In winter, the soil was saturated, and the vegetation was green. Leaf litter was observed in both summer and winter.

#### **King Grass Crop** (5.32482° N and 72.28704° W)

This habitat was a King Grass plantation (*Pennisetum hybridum*). In summer, the grass was dry (yellow) and uncontrolled, sometimes broken, but dense, with a height of between 1 and 2 m and interspersed with thorny shrubs and very few scattered trees (Table 1). In winter, the same pasture was green (1.8 m high), and the soil was wet from rainfall. The search for ticks was conditioned by accessibility and the possibility of placing the bases of the CO_2_ traps among the dense grassland. However, the collection points were distributed broadly over the habitat patches.

#### **Star Grass Paddock** (5.31917 N y 72.29163 W)

This habitat was a paddock with African Star Grass (*Cynodon nlemfuensis*) devoted to cattle breeding (*Bos taurus indicus*). The grass reached a height of 20-40 cm in the paddocks and had some scattered bushes (Table 1). In summer, the grass in the paddock was dry and yellow. The soil was dry and cracked due to the drought. In winter, the grass was green, and the soil partially flooded a few centimeters (<10cm after rain); however, it percolated quickly from one day to the next.

The four habitats’ (Riparian Forest, Cocoa Crop, Star Grass Paddock, King Grass Crop) vegetation’s structural characteristics (Table 1), such as dominant species, light availability, number of strata, depth of leaf litter above the soil, percentage of leaf litter moisture, and dominant soil fraction, were qualified and quantified (details in S2 Appendix) in the points where ticks were sampled.

### Collection of *A. mixtum* specimens in the environment

To define habitat use or non-use, the habitat preference and relative niche width by *A. mixtum* free-living stages, the relative abundance response variable of individuals in each habitat (presence indicating resource use and tolerance to abiotic conditions in the habitat) were used as a proxy for resource selection in the habitat. Based on previous field experiences, it was decided to capture free-living stages of *A. mixtum* through two techniques, CO_2_ traps (dry ice) and white flannel dragging transects (see [52–54]).

#### Use of CO_2_ traps

The details of these traps’ construction before their placement are presented in S3 Appendix. According to vegetation cover, the traps were placed in relatively flat sites and areas, which according to locals, were frequented by domestic and wild animals, and considering their proximity to transited areas (corridors) or roads used by campus workers and students. A minimum number of four CO_2_ traps were placed in each habitat and season (February, August), once per habitat type, and separated by a minimum distance of 5 m. Sampling was conducted between 8:00 am and 1:30 pm, and 2:30 and 4:30 pm, or until all dry ice was consumed. The traps were checked every 30 minutes. The ticks caught on the double-faced tape and those nearby, were collected. Twenty-four traps were placed in the summer and 24 in winter. Another seven traps (55 traps in total) were used to ensure the “non-use” of particular habitats in some tick free-living stages (S1 and S2 Tables). The captured specimens were collected with entomological tweezers, deposited in 50 ml centrifuge tubes with 70% ethanol, and labeled. Later, in the laboratory, the specimens were identified and counted according to genus or species, sex, and stage.

#### White flannel dragging transects

Details of this activity’s outputs and the size of the quantified transects are presented in the S3 Appendix. The sampling was done between 8:00 am and 1:30 pm, and between 2:30 and 4:30 pm, covering several habitats in one day. The transects were separated from each other by at least 5 m. A minimum number of four transects per habitat was performed, once in each season of the year. In the riparian forest habitat, flannel trails were performed when possible, following the paths created by the wildlife and tree base or herbaceous creeping vegetation (<70 cm). The total number of flannel transects was 22 for each time of year, for a total of 44 in the two seasons (S3 and S4 Tables). Two teams of three researchers simultaneously carried out the sampling, using the two capture techniques in each habitat. Each team initially installed the corresponding CO_2_ traps in the four habitats. Flannel dragging was carried out in the defined transects between the time intervals of the trap inspections.

### Taxonomic identification of specimens

Stereoscopy was used to identify the morphological characteristics in adults and some characteristics in nymphs. Light microscopy was used to identify certain morphological characteristics in immature individuals (larvae and nymphs) by mounting specimens between lamina and lamella (subsample of the specimens captured in the habitats and seasons). The taxonomic keys of several authors [55] were used for the genus *Amblyomma* and for the *Amblyomma cajennense* complex [56–59]. The descriptions made by [48] was used to define the species *A. mixtum*.

### Inference of use, habitat preference, and niche width

This study assumed that the presence of ticks in a habitat is possible because: 1) ticks use one or more resources (even if we do not know exactly which or what those resources are, given the lack of biological data on the species [3], and 2) ticks tolerate the habitat’s environmental conditions where we found them [3]. Therefore, we used a paired used/availability habitat design with four discrete levels of habitats with different types of vegetation, such as tropical, riparian evergreen forest, tall grass plantation (King Grass), grazing grass (Star Grass pastures), and cocoa plantation.

We quantified the proportional availability area of a particular habitat. Then, we quantified the observations of use in that habitat at the population level. We performed a graphical analysis with these two variables, comparing the four discrete vegetation types’ proportional use against each habitat’s proportional availability [60]. Using the HaviStat 2.4 free access program [10,61], we explored the visual data to infer relative use and habitat preference between the categories compared (vegetation types, months of different seasons). The greater the disproportion between these two variables in the graph, the higher the preference [8]. Ultimately, a preference was determined when the abundance exceeded the potential habitat area to be used (availability); otherwise, there is use and no preference [2,8]. As stated by [62], “resource selection” can be established when the resource use is disproportionately greater than resource availability; this is assumed correlated with the animal’s fitness [63].

To confirm the graphic approach to habitat use (presence of the species or its stages), habitat preference (asymmetric distribution of resources), or non-use (non-presence of the species in the habitat), five preference indices were calculated using the HaviStat 2.4 program, including Duncan’s index [64], Ivlev’s Electivility Index [65], Bailey confidence interval [66], Alpha index [67,68], constant resource rate (*Constant Resources*) [69], and Interpretation of II ([70]). The HaviStat 2.4 program allows to: 1) check if the sample size is adequate (power, significance level, G-test, Chi-test, and Cherry-test [71]), which is useful for generating inferences at the population level and proposing management strategies [3]; 2) statistically compare field data (95% confidence intervals); and 3) estimate five niche amplitude indices (Standardized Shannon [72], Ivlev’s Electivility Index [65], Levins index [73], Standardized Levins Index [74], and Levins Index Standardized modified by [75]).

The niche width indices were applied to infer whether the habitat resources were used or exploited uniformly (or asymmetrically) by the observed species and the conditions that allow its survival there. In this study, we measured how much wider the spectrum of resources used was in the habitat and how a species could potentially live in a broad range of conditions; this was indicated by its relative abundance (proxy), and the number of habitats inhabited [14]. Regarding the latter, resources and conditions should be considered as a whole, without specific differentiation, which is impossible.

There is not enough published information about *A. mixtum* regarding its life history and autoecology (species-resources-condition-habitat relationships) in the current literature. The Havistat program has been successfully employed in other biological research [8,76–79]. By reaching a consensus among the indices, it is possible to differentiate whether a species uses one habitat or several, and, therefore, its resources), if it prefers that habitat (or others) or strongly selects or avoids one or more particular habitats. According to [62], resources are “avoided” in a habitat when resource use (relative abundance) is disproportionately less than expected, based on the availability of that resource (potentially available habitat). Consequently, even avoided resources may still be used by animals.

Therefore, we speak of a narrow niche for a tick, such as *A. mixtum*, if it only uses or prefers a few hosts to feed and develop, or has little tolerance to environmental variables (e.g., temperature). In the same way, the greater habitat use or preference (e.g., vegetation types) by a tick species (to survive, molt, oviposit, wait for a host and complete the life cycle) and the greater its tolerance to environmental variables, the greater its niche width will be. The previous is derived from specific adaptations to tolerate different conditions and take advantage of heterogeneous or ephemeral resources within each habitat (generalist species [11]).

### Mammal species observed in the four habitats

First, the literature review was used to identify the mammal species most likely to be found in the study area. Posters were made with photographs of these mammal species. They were shown to the people who work, live, or manage the Matepantano farm to identify them photographically (n= 75 people). *In situ* photographic evidence and videos taken by locals, as well as compatible feces, were also used to confirm the species. A list was made of the domestic and wild hosts potentially present on the farm. It was contrasted against the mammal species that, according to the literature, are commonly parasitized by *A. cajennense* s.l. Similarly, qualitative indications were obtained of the relative abundances of host populations for summer and winter. This information was collected for each of the four habitats compared. Moreover, hosts that use to getting closer to the campus buildings and residences were included.

### Climatological analysis of the region

The maximum and minimum range of oscillation of different key environmental variables was calculated for *A. mixtum*, representing its ecological amplitude (niche width). The species’ adaptability to adjust to environmental changes and tolerance level to extreme abiotic values provided the broad spectrum of conditions in the habitats that allow it to survive, complete the life cycle, and persist. To this end, a 29-year analysis was made of the climatological data available (1975-2004) from two of the Colombian Institute of Hydrology, Meteorology, and Environmental Studies’ (IDEAM in Spanish) meteorological stations, the IDEAM, main automatic telemetry weather station at the Yopal airport - code 35215020- and the Yopal conventional pluviometric station-code 35210020. The consultation was made in 2019. Both stations are located in Yopal (03.05 N; 76.33 W; 1,205 m.a.s.l.), which is within a 20 km radius from the Matepantano farm. This information was complemented with the multi-year analysis (1981-2010) provided by the IDEAM (available at http://atlas.ideam.gov.co/visorAtlasClimatologico.html). Because of its spatial scale, we termed this climate analysis “regional level.”

### Local weather and microclimate information

The regional information was complemented with data from a permanent station for continuous climate recording located in the cocoa crop sampled at the Matepantano farm. This station, called Davis, has registered for seven years (2012 to 2019, every 30 minutes) temperature, relative humidity, solar radiation, solar energy, rainfall, and wind speed using a Davis-Vantage Pro2 unit located at an altitude of 1.2 m and 300 m from the Matepantano farm’s main buildings. We also calculated the range of oscillation (maximum and minimum value) of different key environmental variables for the tick (ecological amplitude of *A. mixtum*). Because of its spatial scale, we termed this climate analysis “local level.”

To determine the range of daily oscillation experienced by ticks at the microhabitat level, the temperature, humidity, and solar irradiance were measured *in situ* in the Cocoa Crop, Star Grass Paddock, and riparian forest in summer (February) and in the Cocoa Crop and King Grass Crop in winter (August). Hobo S-THB-M8000 sensors were used for temperature (accuracy ±0,2°C) and relative humidity (accuracy ±2,5%) and the Hobo S-LIB-MOO3 sensor for solar radiation (irradiance) (accuracy ±10 W/m^2^). The measurements were taken 30 cm from the ground, every 30 seconds in summer and every minute in winter. In summer, the variables were measured the morning (9.30 am to 12 pm) of February 8^th^, 2019, in the cacao plantation, the afternoon (2.30 to 4.30 pm) of February 8^th^ and 9^th^ in the Star Grass paddock, and all day (9.30 am to 4.30 pm) on February 10^th^ in the riparian forest. In winter, on August 16^th^, in the cocoa plantation and in King Grass crop on August 17^th^, the same three variables were measured in the morning and/or the afternoon. The three climatic variables were measured in winter at 30 cm above the ground and only for cocoa crop habitat within the leaf litter (a refuge usually frequented by ticks).

## RESULTS

### Overview of the relative abundance of stages of *A. mixtum*

The collected and taxonomically classified *Amblyomma* ticks corresponded to the species *A. mixtum*. Nymphs dominated the relative abundance of *A. mixtum* in the summer season (February) for all selected habitats with 59.7% (4,021/6,733), larvae had 37.6% (2,533/6,733), and adults a low 2.7% (179/6,733). Meanwhile, in winter (August), the captured population not only decreased by one order of magnitude but was dominated by adults with 61.7% (164/266), in three of the four habitats except the Star Grass paddock, followed by larvae with 30.1% (80/266), and with few nymphs 8.3% (22/266) (Table 2).

**Table 2.** Number of collected and identified *A. mixtum* specimens according to collection method, habitat type and stage.

|  |
| --- |
| (A) Collection carried out on February, 2019 (summer) |

| Collection method | Capture fraction* | Habitat | Larvae | Nymphs | Females | Males | TOTAL |
| --- | --- | --- | --- | --- | --- | --- | --- |
| CO<sub>2</sub> traps | 7/7 | Star Grass Paddock | 205 | 431 | 17 | 3 | 656 |
|  | 5/5 | King Grass Crop | 212 | 1,882 | 22 | 52 | 2,168 |
|  | 8/8 | Riparian Forest | 532 | 757 | 39 | 24 | 1,352 |
|  | 4/4 | Cocoa Crop | 61 | 729 | 10 | 5 | 805 |
| **Subtotal** |  |  | **1,010** | **3,799** | **88** | **84** | **4,981** |
| Cloth dragging by transects | 7/7 | Star Grass Paddock | 198 | 37 | 0 | 0 | 235 |
|  | 6/6 | King Grass Crop | 496 | 74 | 0 | 0 | 570 |
|  | 4/4 | Riparian Forest | 214 | 38 | 2 | 0 | 254 |
|  | 5/5 | Cocoa Crop | 615 | 73 | 4 | 1 | 693 |
| **Subtotal** |  |  | **1,523** | **222** | **6** | **1** | **1,752** |
| **TOTAL** |  |  | **2,533** | **4,021** | **94** | **85** | **6,733** |
| **(B) Collection carried out on August, 2019 (Winter)** |  |  |  |  |  |  |  |
| Collection method | Capture fraction* | Habitat | Larvae | Nymphs | Females | Males | TOTAL |
| CO<sub>2</sub> traps | 2/8 | Star Grass Paddock | 0 | 10 | 12 | 10 | 32 |
|  | 6/6 | King Grass Crop | 0 | 4 | 28 | 27 | 59 |
|  | 11/12 | Riparian Forest | 0 | 5 | 18 | 26 | 49 |
|  | 5/5 | Cocoa Crop | 0 | 0 | 13 | 22 | 35 |
| **Subtotal** |  |  | **0** | **19** | **71** | **85** | **175** |
| Cloth dragging by transects | 2/5 | Star Grass Paddock | 67 | 0 | 0 | 0 | 67 |
|  | 2/7 | King Grass Crop | 13 | 0 | 1 | 0 | 14 |
|  | 3/4 | Riparian Forest | 0 | 3 | 4 | 1 | 8 |
|  | 2/6 | Cocoa Crop | 0 | 0 | 1 | 1 | 2 |
| **Subtotal** |  |  | **80** | **3** | **6** | **2** | **91** |
| **TOTAL** |  |  | **80** | **22** | **77** | **87** | **266** |

(A) summer; (B) winter at thr Yopal (Casanare).
\*Number of traps or transects with effective capture related to the total number of traps or transects that were located in each habitat.

*A. mixtum*’s three stages (larvae, nymphs, and adults) used all four habitats with different proportions between the two periods of the year studied (Fig 3A and 3B and Table 2). In summer, 40% (2,738/6,733) of *A. mixtum* ticks were found in the King Grass Crop, 23.9% (1,606/6,733) in the Riparian Forest, 22.2% (1,498/6,733) in the Cocoa Crop, and 13.2% (891/6,733) in the Star Grass Paddock (Table 2). In winter, 37.2% (99/266) of the ticks used the Star Grass Paddock, followed by 27.4% (73/266) in the King Grass plantation, 21.4% (57/266) in the Riparian Forest, and 13.9% (37/266) in the Cocoa Crop (Table 2).

**Figure 3.**
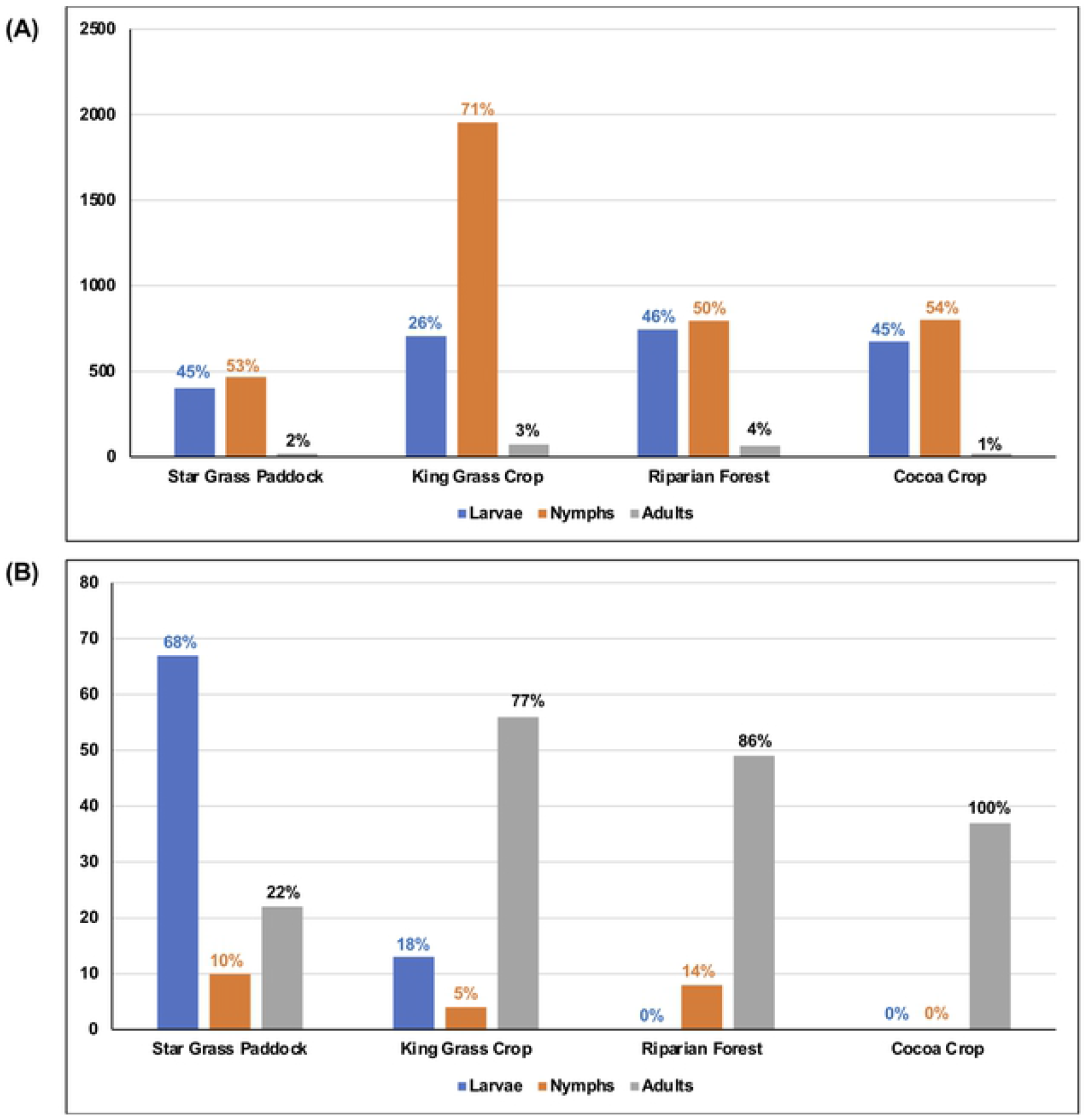
Abundance of *A. mixtum* in every sampled habitat according to the season in the Matepantano farm. (A). February of 2019 (summer). (B). August of 2019 (winter).

In summer, the three stages of *A. mixtum* used all the habitats (Figs 3A and 4A, and Tables 2–5). Only the adult stage of *A. mixtum* used all the habitats in summer and winter. Its absolute abundance remained relatively similar in both seasons. In the winter season, the nymphs did not use the Cocoa Crop habitat, and the larvae were absent in the Riparian Forest and Cocoa Crop habitats (Figures 3B and 4B, and Tables 2–5). *A. mixtum*’s gender ratio for the total adults in both periods (females:males with 171:172) was 0.994.

**Figure 4.**
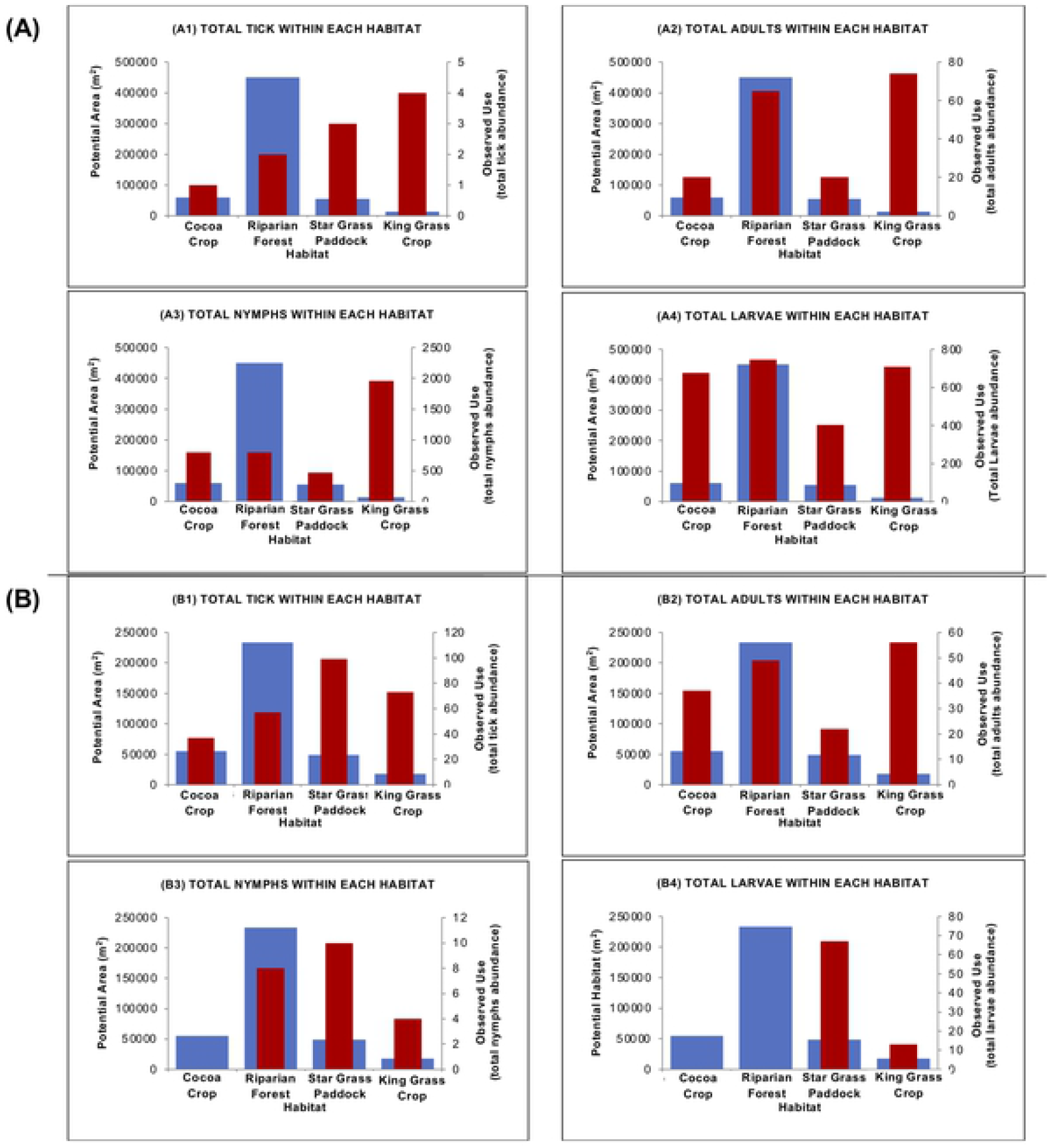
Trend graphs to infer the use and habitat preference of *A. mixtum*. These plots relate tick total abundance and tick stage abundance of both seasons, summer and winter, (as an equivalent of observed use of each habitat resources) (red bar 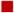; frequency on the y’-axis on the right) and the potential or available area of the habitat that *A. mixtum* might use (blue bar 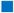; frequency on the y-axis on the left) at the Matepanano farm. In that way, the higher the red bar and the lower the blue bar, the greater the habitat preference. When the red bar barely exceed or equals the blue bar, the relationship is correspond to habitat usage. When the blue bar far surpasses the red bar, there is avoidance of such a habitat (see tables 3–5). (A). Summer season. (A1) The mathematical index points out that *A. mixtum* uses all four habitats. In addition, *A. mixtum* population (stage independent) uses three habitats and shows preference for ‘Star Grass Paddock’ and ‘King Grass Crop’, while it avoids the ‘Riparian Forest’ habitat. (A2) Adults of *A. mixtum* prefer ‘King Grass Crop’, while they use the remainder habitats. (A3) Nymphs of *A. mixtum* prefer ‘King Grass Crop’ and ‘ Cocoa Crop’, they use ‘Star Grass Paddock’, but they avoid the ‘Riparian Forest’ habitat. (A4) Larvae of *A. mixtum* prefers three habitats (‘King Grass Crop’, ‘Cocoa Crop’ and ‘Star Grass Paddock’), while they only use the ‘Riparian Forest’ habitat. (B). Winter season. (B1) *A. mixtum* population (stage independent) prefers ‘King Grass Crop’ and ‘Star Grass Paddock’, it uses ‘Cocoa Crop’ and avoids ‘Riparian Forest’. (B2) Adults of *A. mixtum* prefer ‘King Grass Crop’ and ‘Cacao Crop’, while they use ‘Star Grass Paddock’ and ‘Riparian Forest’. (B3) Nymphs of *A. mixtum* uses ‘Cocoa Crop’, the prefer ‘Star Grass Paddock’ and ‘King Grass Crop’, and they avoids the ‘Riparian Forest’ habitat. (B4) Larvae of *A. mixtum* only uses two out of four habitats, preferring ‘Star Grass Paddock’ and using ‘King Grass Crop’.

In total, 2.8 times more ticks were caught with CO_2_ traps (4,981) than with white flannelettes transects (1,752) in the summer time. In winter, the CO_2_ traps’ effectiveness for tick collection decreased 1.9 times (175 individuals) compared to the transects (91 individuals). The flannel dragging method in the pre-determined transects was most effective in capturing the larvae in summer and winter compared to the remnant stages (Table 2). Up to 17 times more nymphs and 25 times more adults were collected using the CO_2_ traps than in the transects in summer. Similarly, six times more nymphs and 20 times more adults were captured in winter using CO_2_ traps than flannel dragging. In all habitats, CO_2_ traps were more effective in collecting ticks in summer; in winter, they were effective in 75% of the habitats. The transects only outperformed the traps in capturing ticks in the Star Grass paddock habitat in winter (Table 2).

### Habitat use and preference

The Cherry Test [71] indicated that in summer and winter the total sample of *A. mixtum* ticks captured was sufficient to infer on the use and preference (Table 3). It should be noted that, during the winter season, nymphs were absent from sampling in the cocoa plantation habitat and larvae were not found in the cocoa plantation and riparian forest habitats. Duncan’s Preference Indexes [64], of Ivlev’s electivity [65], Bailey’s confidence interval [66], Alpha [67,68] (Constant Resources [69]), and Interpretation of II [70,74] indicated that, the *A. mixtum* population used all four habitats (Figure 4 and Table 4). Moreover, most of these indices indicated *A. mixtum*’s preference for three of the four habitats (King Grass Crop, Cocoa Crop, and Star Grass Paddock).

**Table 3.**
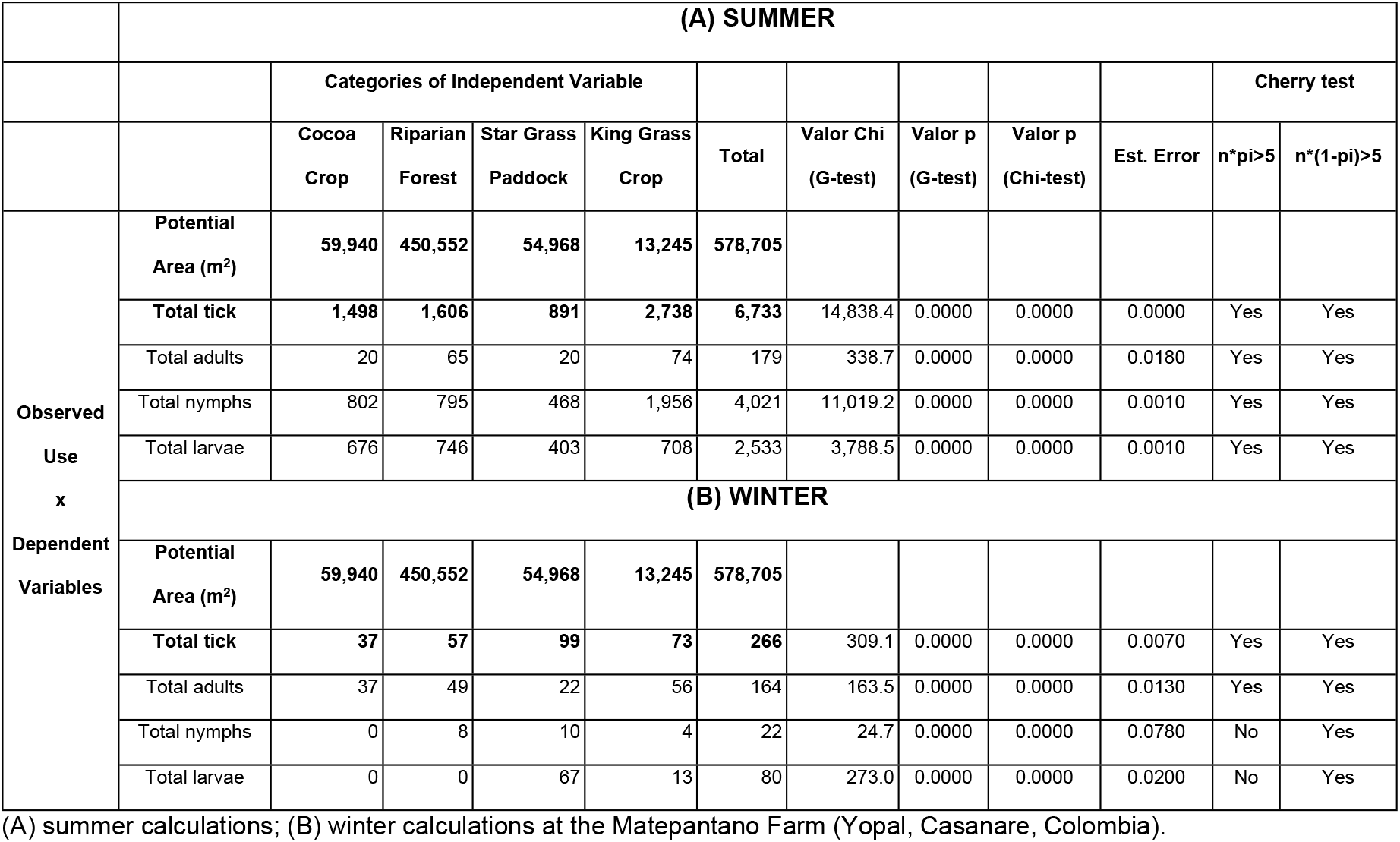
Calculations of potential area and *A. mixtum* abundance within each habitat to obtain the observed use.

**Table 4.** Calculations of the best five habitat preferences indices for *A. mixtum* in the Matepantano Farm.

| (A) SUMMER |  |  |  |  |  |
| --- | --- | --- | --- | --- | --- |
| HABITAT PREFERENCE INDICES |  |  |  |  | NOTES |
| DUNCAN INDEX [64] |  |  |  |  |  |
| Categories of Independent Variable |  |  |  |  |  |
|  | Cocoa Crop | Riparian Forest | Star Grass Paddock | King Grass Crop |  |
| Total tick abundance | 0.5 | 0.1 | 0.4 | 1.3 | Range: 0 to +Inf.<br>If > 0.3 = Preference<br>If < 0.3 = No Preference |
| Total adults | 0.3 | 0.2 | 0.3 | 1.3 |  |
| Total nymph | 0.5 | 0.1 | 0.4 | 1.3 |  |
| Total larvae | 0.6 | 0.1 | 0.4 | 1.1 |  |
| IVLEV'S ELECTIVITY INDEX [65] |  |  |  |  |  |
| Categories of Independent Variable |  |  |  |  |  |
|  | Cocoa Crop | Riparian Forest | Star Grass Paddock | King Grass Crop |  |
| Total tick abundance | 0.4 | -0.5 | 0.2 | 0.9 | Range: -1 to +1<br>If > 0 = Preference<br>If < 0 = No Preference |
| Total adults | 0.0 | -0.4 | 0.1 | 0.9 |  |
| Total nymph | 0.3 | -0.6 | 0.1 | 0.9 |  |
| Total larvae | 0.4 | -0.5 | 0.3 | 0.8 |  |
|  | **BAILEY CONFIDENCE INTERVALS [66]** |  |  |  |  |
|  | **Categories of Independent Variable** |  |  |  |  |
|  | **Cocoa Crop** | **Riparian Forest** | **Star Grass Paddock** | **King Grass Crop** |  |
| **Total tick abundance** | **697.4** | 5,242.0 | **639.5** | **154.1** | **If upper Interval < Species Use = Preference**<br>If Lower interval > Species Use = No Preference<br>If Lower Interval < Species Use < Upper Interval = Use |
| **Total adults** | 18.5 | 139.4 | 17.0 | **4.1** |  |
| **Total nymph** | **416.5** | 3,130.6 | **381.9** | **92.0** |  |
| **Total larvae** | **262.4** | 1,972.1 | **240.6** | **58.0** |  |
|  | **ALPHA INDEX [67,68] (CONSTANT RESOURCES [69])** |  |  |  |  |
|  | **Categories of Independent Variable** |  |  |  |  |
|  | **Cocoa Crop** | **Riparian Forest** | **Star Grass Paddock** | **King Grass Crop** |  |
| **Total tick abundance** | 0.1 | 0.0 | 0.1 | **0.8** | **Range: 0 to +1** |
| **Total adults** | 0.1 | 0.0 | 0.1 | **0.9** | **If > 1/(#Indep. Var.) = Preference** |
| **Total nymph** | 0.1 | 0.0 | 0.0 | **0.9** | If < 1/(#Indep. Var.) = No Preference |
| **Total larvae** | 0.2 | 0.0 | 0.1 | **0.7** | 1/(#Indep. Var.) = 0.25 |
|  | **INTERPRETATION OF II [70]** |  |  |  |  |
|  | **Categories of Independent Variable** |  |  |  |  |
|  | **Cocoa Crop** | **Riparian Forest** | **Star Grass Paddock** | **King Grass Crop** | **Range: -1 to +1** |
| **Total tick abundance** | 0.4 | -0.8 | 0.2 | **0.9** | If -1 < Index Value < -0.5 = Strong Avoidance |
| **Total adults** | 0.0 | -0.7 | 0.1 | **0.9** | If -0.49 < Index Value < -0.26 = Moderate Avoidance |
| **Total nymph** | 0.4 | -0.9 | 0.1 | **1.0** | If -0.25 < Index Value < 0.25 = Indifference |
| **Total larvae** | 0.5 | -0.8 | 0.3 | **0.9** | If 0.26 < Index Value < 0.49 = Neutral Selection |
|  |  |  |  |  | **If 0.50 < Index Value < 1 = Strong Selection** |
| **(B) WINTER** |  |  |  |  |  |
|  | **HABITAT PREFERENCE INDICES** |  |  |  | **NOTES** |
|  | **DUNCAN INDEX [64]** |  |  |  |  |
|  | **Categories of Independent Variable** |  |  |  |  |
|  | **Cocoa Crop** | **Riparian Forest** | **Star Grass Paddock** | **King Grass Crop** |  |
| **Total tick abundance** | 0.3 | 0.1 | **0.6** | **0.8** | **Range: 0 to +Inf.** |
| **Total adults** | **0.4** | 0.2 | 0.3 | **0.9** | **If > 0.3 = Preference** |
| **Total nymph** | - | 0.2 | **0.6** | **0.7** | If < 0.3 = No Preference |
| **Total larvae** | - | - | **0.9** | **0.6** | - = absence of individuals |
|  | **IVLEV'S ELECTIVITY INDEX [65]** |  |  |  |  |
|  | **Categories of Independent Variable** |  |  |  |  |
|  | **Cocoa Crop** | **Riparian Forest** | **Star Grass Paddock** | **King Grass Crop** |  |
| **Total tick abundance** | -0.1 | -0.5 | **0.5** | **0.7** | **Range: -1 to +1** |
| **Total adults** | **0.2** | -0.4 | 0.0 | **0.7** | **If > 0 = Preference** |
| **Total nymph** | - | -0.3 | **0.5** | **0.6** | If < 0 = No Preference |
| **Total larvae** | - | - | **0.7** | **0.5** | - = absence of individuals |
|  | **BAILEY CONFIDENCE INTERVALS [66]** |  |  |  |  |
|  | **Categories of Independent Variable** |  |  |  |  |

|  | Cocoa Crop | Riparian Forest | Star Grass Paddock | King Grass Crop |  |
| --- | --- | --- | --- | --- | --- |
| Total tick abundance | 41.1 | 175.0 | 36.4 | 13.5 | **If upper Interval < Species Use = Preference**<br>If Lower interval > Species Use = No Preference<br>If Lower Interval < Species Use < Upper Interval = Use<br>- = absence of individuals |
| Total adults | 25.4 | 107.9 | 22.5 | 8.3 |  |
| Total nymph | - | 14.5 | 3.0 | 1.1 |  |
| Total larvae | - | - | 11.0 | 4.1 |  |
| ALPHA INDEX [67,68] (CONSTANT RESOURCES [69]) |  |  |  |  |  |
| Categories of Independent Variable |  |  |  |  |  |
|  | Cocoa Crop | Riparian Forest | Star Grass Paddock | King Grass Crop | Range: 0 to +1<br>**If > 1/(#Independent Variable) = Preference**<br>If < 1/(#Independent Variable) = No Preference<br>1/(#Independent Variable) = 0.25<br>- = absence of individuals |
| Total tick abundance | 0.1 | 0.0 | 0.3 | 0.6 |  |
| Total adults | 0.2 | 0.0 | 0.1 | 0.7 |  |
| Total nymph | - | 0.1 | 0.4 | 0.5 |  |
| Total larvae | - | - | 0.7 | 0.3 |  |
| INTERPRETATION OF II [70] |  |  |  |  |  |
|  |  |  |  |  | Range: -1 to +1 |
| Categories of Independent Variable |  |  |  |  | If -1 < Index Value < -0.5 = Strong Avoidance |
|  | Cocoa Crop | Riparian Forest | Star Grass Paddock | King Grass Crop | If -0.49 < Index Value < -0.26 = Moderate Avoidance |
| Total tick abundance | -0.1 | -0.8 | 0.6 | 0.8 | If -0.25 < Index Value < 0.25 = Indifference |
| Total adults | 0.2 | -0.6 | 0.0 | 0.8 | If 0.26 < Index Value < 0.49 = Neutral Selection |
| Total nymph | - | -0.5 | 0.7 | 0.6 | **If 0.50 < Index Value < 1 = Strong Selection** |
| Total larvae | - | - | 0.9 | 0.6 | - = absence of individuals |

*A. mixtum* uses, but does not prefer, the Riparian Forest habitat. This result was consistent for nymphs, larvae, and adults, which was corroborated by the interpretation of II index that indicated a strong avoidance of this habitat (Table 5). Although adults also preferred the King Grass plantation habitat in both seasons, according to all selected indices, in summer, they used, but did not prefer the cocoa plantation and Star Grass Paddock, according to four of the five indices (Table 4). In winter, the concordance of non-preference of adults decreases between indices with respect to the habitat ‘Cocoa Crop’ but remains with respect to ‘Star Grass Paddock’. The interpretation index of II shows a strong avoidance of the habitat ‘Riparian Forest ‘ by the adults (Table 4). In the case of larvae and nymphs in summer the concordance of preference between indices is not complete either. Larvae and nymphs prefer three of the four habitats according to three of the five selected indices. In winter the five indices coincided in indicating preference of the immature stages for the habitats ‘Star Grass Paddock’ and ‘ King Grass Crop’ (Table 4).

**Table 5.** Calculations of the best five niche amplitud indices for *A. mixtum* within its habitats in the Matepantano Farm.

| (A) SUMMER |  |  |  |  |  |  |
| --- | --- | --- | --- | --- | --- | --- |
|  | NICHE AMPLITUDE INDICES |  |  |  |  | NOTES |
|  | STANDARDIZED SHANNON INDEX [72] |  |  |  |  |  |
|  | Categories of Independent Variable |  |  |  |  |  |
|  | Cocoa Crop | Riparian Forest | Star Grass Paddock | King Grass Crop | Index | Bolded value = maximum amplitude |
| Potential Area (m²) | 59,940 | 450,552 | 54,968 | 13,245 | Value | *Italic value* = minimum amplitude |
| Total tick abundance | 1,498 | 1,606 | 891 | 2,738 | 0.94 | Range: 0 to +1<br>0 = minimum niche amplitude<br>+1 =maximum niche amplitude |
| Total adults | 20 | 65 | 20 | 74 | 0.88 |  |
| Total nymphs | 802 | 795 | 468 | 1,956 | 0.90 |  |
| Total larvae | 676 | 746 | 403 | 708 | 0.98 |  |
|  | IVLEV'S ELECTIVITY INDEX [65] |  |  |  |  |  |
|  | Categories of Independent Variable |  |  |  |  |  |
|  | Cocoa Crop | Riparian Forest | Star Grass Paddock | King Grass Crop |  |  |
| Potential Area (m²) | 59,940 | 450,552 | 54,968 | 13,245 |  |  |
| Total tick abundance | 1,498 | 1,606 | 891 | 2,738 | 1.95 | Range: 0 to +infinite<br>0 = maximum niche amplitude<br>+infinite = minimum niche amplitude |
| Total adults | 20 | 65 | 20 | 74 | 1.38 |  |
| Total nymphs | 802 | 795 | 468 | 1,956 | 1.92 |  |
| Total larvae | 676 | 746 | 403 | 708 | 1.99 |  |
|  | LEVINS INDEX [73] |  |  |  |  |  |
|  | Categories of Independent Variable |  |  |  |  |  |
|  | Cocoa Crop | Riparian Forest | Star Grass Paddock | King Grass Crop |  |  |
| Potential Area (m²) | 59,940 | 450,552 | 54,968 | 13,245 |  |  |
| Total tick abundance | 1,498 | 1,606 | 891 | 2,738 | 1.31 | Range: 0 to +2<br>0 = minimum niche amplitude<br>+2 = maximum niche amplitude |
| Total adults | 20 | 65 | 20 | 74 | 1.22 |  |
| Total nymphs | 802 | 795 | 468 | 1,956 | 1.24 |  |
| Total larvae | 676 | 746 | 403 | 708 | 1.36 |  |
|  | STANDARDIZED LEVINS INDEX [74] |  |  |  |  |  |
|  | Categories of Independent Variable |  |  |  |  |  |
|  | Cocoa Crop | Riparian Forest | Star Grass Paddock | King Grass Crop |  |  |
| Potential Area (m²) | 59,940 | 450,552 | 54,968 | 13,245 |  |  |
| Total tick abundance | 1,498 | 1,606 | 891 | 2,738 | 0.82 | Range: 0 to +1<br>0 = minimum niche amplitude<br>+1 = maximum niche amplitude |
| Total adults | 20 | 65 | 20 | 74 | 0.68 |  |
| Total nymphs | 802 | 795 | 468 | 1,956 | 0.68 |  |
| Total larvae | 676 | 746 | 403 | 708 | 0.94 |  |
|  | MODIFIED STANDARDIZED LEVINS INDEX [75] |  |  |  |  |  |
|  | Categories of Independent Variable |  |  |  |  |  |
|  | Cocoa Crop | Riparian Forest | Star Grass Paddock | King Grass Crop |  |  |
| Potential Area (m²) | 59,940 | 450,552 | 54,968 | 13,245 |  |  |
| Total tick abundance | 1,498 | 1,606 | 891 | 2,738 | 0.86 | Range: (1/Independent Variable) to +1<br>(1/Independent Variable) = minimum<br>+1 = maximum niche amplitude |
| Total adults | 20 | 65 | 20 | 74 | 0.76 |  |
| Total nymphs | 802 | 795 | 468 | 1,956 | 0.76 |  |
| Total larvae | 676 | 746 | 403 | 708 | 0.96 |  |
| (B) WINTER |  |  |  |  |  |  |
|  | NICHE AMPLITUDE INDICES |  |  |  |  | NOTES |
|  | STANDARDIZED SHANNON INDEX [72] |  |  |  |  |  |
|  | Categories of Independent Variable |  |  |  |  |  |
|  | Cocoa Crop | Riparian Forest | Star Grass Paddock | King Grass Crop | Index | Bolded value = maximum amplitude |
| Potential Area (m²) | 59,940 | 450,552 | 54,968 | 13,245 | Value | Italicize value = minimum amplitude |
| Total tick abundance | 37 | 57 | 99 | 73 | 0.96 | Range: 0 to +1<br>0 = minimum niche amplitude<br>+1 =maximum niche amplitude<br>- = absence of individuals |
| Total adults | 37 | 49 | 22 | 56 | 0.96 |  |
| Total nymphs | - | 8 | 10 | 4 | 0.75 |  |
| Total larvae | - | - | 67 | 13 | 0.32 |  |
|  | IVLEV'S ELECTIVITY INDEX [65] |  |  |  |  |  |
|  | Categories of Independent Variable |  |  |  |  |  |
|  | Cocoa Crop | Riparian Forest | Star Grass Paddock | King Grass Crop |  |  |
| Potential Area (m²) | 59,940 | 450,552 | 54,968 | 13,245 |  |  |
| Total tick abundance | 37 | 57 | 99 | 73 | 1.71 | Range: 0 to +infinite<br>0 = maximum niche amplitude<br>+infinite = minimum niche amplitude |
| Total adults | 37 | 49 | 22 | 56 | 1.31 |  |
| Total nymphs | - | 8 | 10 | 4 | 2.39 |  |
| Total larvae | - | - | 67 | 13 | 3.24 | - = absence of individuals |
| LEVINS INDEX [73] |  |  |  |  |  |  |
| Categories of Independent Variable |  |  |  |  |  |  |
|  | Cocoa Crop | Riparian Forest | Star Grass Paddock | King Grass Crop |  |  |
| Potential Area (m<sup>2</sup>) | 59,940 | 450,552 | 54,968 | 13,245 |  |  |
| Total tick abundance | 37 | 57 | 99 | 73 | 1.33 | Range: 0 to +2 |
| Total adults | 37 | 49 | 22 | 56 | 1.33 | 0 = minimum niche amplitude |
| Total nymphs | - | 8 | 10 | 4 | 1.04 | +2 = maximum niche amplitude |
| Total larvae | - | - | 67 | 13 | 0.44 | - = absence of individuals |
| STANDARDIZED LEVINS INDEX [74] |  |  |  |  |  |  |
| Categories of Independent Variable |  |  |  |  |  |  |
|  | Cocoa Crop | Riparian Forest | Star Grass Paddock | King Grass Crop |  |  |
| Potential Area (m<sup>2</sup>) | 59,940 | 450,552 | 54,968 | 13,245 |  |  |
| Total tick abundance | 37 | 57 | 99 | 73 | 0.86 | Range: 0 to +1 |
| Total adults | 37 | 49 | 22 | 56 | 0.88 | 0 = minimum niche amplitude |
| Total nymphs | - | 8 | 10 | 4 | 0.56 | +1 = maximum niche amplitude |
| Total larvae | - | - | 67 | 13 | 0.12 | - = absence of individuals |
| MODIFIED STANDARDIZED LEVINS INDEX [75] |  |  |  |  |  |  |
| Categories of Independent Variable |  |  |  |  |  |  |
|  | Cocoa Crop | Riparian Forest | Star Grass Paddock | King Grass Crop |  |  |
| Potential Area (m<sup>2</sup>) | 59,940 | 450,552 | 54,968 | 13,245 |  |  |
| Total tick abundance | 37 | 57 | 99 | 73 | 0.90 | Range: (1/Independent Variable) to +1 |
| Total adults | 37 | 49 | 22 | 56 | 0.91 | (1/Independent Variable) = minimum |
| Total nymphs | - | 8 | 10 | 4 | 0.67 | +1 = maximum niche amplitude |
| Total larvae | - | - | 67 | 13 | 0.34 | - = absence of individuals |

(A) summer calculations; (B) winter calculations using HaviStat 2.4®.

### Niche width

In summer, the niche amplitude was maximum for the larvae in four of the five indices, as the value of Ivlev’s electivity index was the lowest. In general, the rates of niche width obtained in this part of the year are considered high for all free-living stages of *A. mixtum*, as well as the population as a whole, since the values obtained tend towards the maximum amplitude (Table 5) [65,72,73,75]. Interestingly, during the winter season, the five selected indices coincided in a maximum niche amplitude for the adult stage (Table 5), which also coincides with similar abundance data for adults in the two sampled seasons (Table 2). That is, in summer and winter the survival rate of adults was similar and could indicate that adults are present at different times throughout the year. In contrast, the population of larvae and nymphs almost succumbed in winter, reducing their numbers by one order of magnitude when compared to their collection in summer and disappearing from the ‘Cocoa Plantation’ habitat. The larvae were also not found in the ‘Riparian Forest’ habitat (Figure 3 and Table 2).

Remarkably, the larvae in winter obtained the lowest niche amplitude in all the selected indices (Table 5). This suggests that adults and immature *A. mixtum* would have differentiated tolerance to abiotic conditions and different biotic relationships or behaviors in the studied habitats. Events derived from winter weather behavior (e.g., prolonged flooding; favorable conditions for the development of natural enemies such as fungi; scarcity of resources) could configure a greater vulnerability for larvae and nymphs in ‘Cocoa Crop’, ‘Riparian Forest’ and ‘King Grass Crop’ (Figure 3). Therefore, the seasonality observed at the Matepantano farm, and its climatic and environmental variability, may affect the use, preference and extent of the niche of the free-living stages of *A. mixtum* that, which respond differently to the habitat.

### Climate of the region in summer and winter

Wide seasonal variation of ambient temperature is observed in the region of up to 12 °C on average between the summer and winter seasons (Table 6) and an absolute range of 26°C at the local weather station located in the ‘Cocoa Plantation’ habitat (Table 7). The temperature variation in summer at the Matepantano farm is wider (from 18°C to 37°C) when compared to the winter season (minimum of 19°C and maximum of 33.6°C). Total precipitation in winter (August) was one order of magnitude higher than in summer (Table 7). The average relative humidity varied by 20% between February and August, both in the regional (61 and 81%) and local (68 and 88%) climate seasons, reaching minimum values of 30% in the summer season (Tables 6 and 7).

**Table 6.** Multiannual mean range for several climatic variables recorded during 29 years (1975-2004) in two weather stations located at the Yopal municipality (Casanare, Colombia).

| Variable | Range of the multiannual mean* | Observations |
| --- | --- | --- |
| Temperature (°C) | 24-28 | The driest period was from November to March. Maximum and minimum values of the multiannual mean temperature during the 29 years period were 33,7 and 30,1 °C for February, and 23,8 and 21,7 °C for August. |
| Total precipitation (mm) | 2.500-3.000 | An unimodal rainfall cycle with a peak from April to October. The maximum precipitation/day reached 200 mm in just one day. The multiannual mean precipitation was 16 mm for February and 97 mm for August. |
| Evaporation (mm/year) | 1.300-1.500 | Almost a half of the rainfall is evaporated. |
| Potential evapotranspiration (mm/year) | 1.200-1.400 |  |
| Relative humidity (%) | 75-80 | Perhumid climatic type (Thornthwaite classification Limit 7/9). Multiannual mean relative humidity between 1981 to 2010 was 61% for February and 81% for August. |
| Daily hours of sunlighth | 5-6 |  |
| Daily solar irradiation (KW/h/m<sup>2</sup>) | 4,5-5 |  |
| Ultraviolet radiation (IUV) | 7-8 |  |
| Soil net water (mm) | 1.000-2.000 | Soil drought is common between the November to January period. |
| Water excess in the annual extreme (decades) | 15-20 |  |
| Water deficit in the annual extreme (decades) | 5-10 |  |
| Wind speed (m/s) | 3-4 | The prevailing wind direction at the study site was East-West. |

\*Data delivered by IDEAM (IDEAM, [Climatic Atlas of Colombia, Interactive version, Bogotá, D.C., 2015, text in Spanish]
ISBN: Volume 1: 978-958-8067-73-5) corresponded to annual mean with annotations about variable behavior in every month.

**Table 7.** Multiannual mean of climatic variables recorded during seven years (2012-2019) in the Matepantano farm.

| Variable* | Mean (SD) | Max. | Min. | Difference | Observations |
| --- | --- | --- | --- | --- | --- |
| Annual temperature (°C) | 25,9(±3,5) | 41,9 | 15,9 | 26,0 | Maximum value was recorded in 2018, while the minimum value was recorded in 2015. |
| Temperatura (°C) in February (summer) | 27,7(±4,1) | 37,0 | 18,0 | 19,0 |  |
| Temperatura (°C) in August (winter) | 25,2(±3,0) | 33,6 | 19,0 | 14,6 |  |
| Total annual precipitation (mm) | 1559,8(±507,7) | 2251 | 619 | 1632 | Maximum precipitation (155 mm in one day) between July to November of 2016. |
| Total precipitation (mm) in February | 23,7(±37,7) | 105 | 0 | 105 |  |
| Total precipitation (mm) in August | 366(±104,3) | 481 | 193 | 288 | It represents almost the third of the annual total precipitation. |
| Relative humidity (%) | 83,4(±13,6) | 100 | 30 | 70 |  |
| Relative humidity (%) in February | 68,0(±14,5) | 99 | 30 | 69 |  |
| Relative humidity (%) in August | 88,3(±8,6) | 100 | 50 | 50 |  |
| Dew point (°C) | 22,2(±4,7) | 27,2 | 18,2 | 9 |  |
| Solar irradiation (KW/m<sup>2</sup>) | 221,7(±317,0) | 1447 | 0 | 1447 |  |
| Solar energy -radiant light and heat- (Exajoules) | 3,3(±6,3) | 46,6 | 0 | 46,6 |  |
| Atmospheric pressure (milibar) | 1.010(±2) |  |  |  |  |
| Wind speed (m/s) | 0,76($\pm 0,36$) | 16,1 | 0 | 16,1 | Often strong gust of wind in the N-NE and E directions.<br>The strongest gust of wind between August to November. Only throughout the 2015 year, the wind direction was to the North. |

\*Data from the weather station located at the 'Cacao Crop' habitat (N = 242,152 records).
**NOTICE:** A milimeter of water equals the emptying of one (1) liter of water in a 1 m<sup>2</sup> surface of enclosed space (the water volume will have exactly 1 mm of thickness). So, the multiannual mean precipitation in February (summer) in the Matepantano farm is 23.7 litres/m<sup>2</sup> that is equal to 23.7 mm or around 2 cm of water over the ground. By contrast, the multiannual mean precipitation in August (winter) is 366 litres/m<sup>2</sup> (366 mm or almost 36 cm of water over the ground). It means around 15 times more ponding per m<sup>2</sup> in August than in February.

### Microclimatic variables by habitat in summer

#### Cocoa Crop

Solar radiation remained low and relatively constant (<200 W/m^2^) in the morning and up to noon (Figure 5), with some very irregular readings (200–800 W/m^2^) probably due to the sustained movement of the wind on the cocoa leaves that allowed intermittent, but direct, exposure to sunlight (Figure 5). In the afternoon, the solar radiation went from almost 500 W/m^2^ up to approximately 200 W/m^2^. Thus, the solar radiation can have variations of up to 400 W/m^2^ in a few hours in the same day. The ambient temperature increased by 9°C (from 25 to about 34°C) from early morning to midday. In the afternoon, the temperature drops from 38°C to 35°C. That is, the temperature can vary up to 13°C in seven hours.

**Figure 5.**
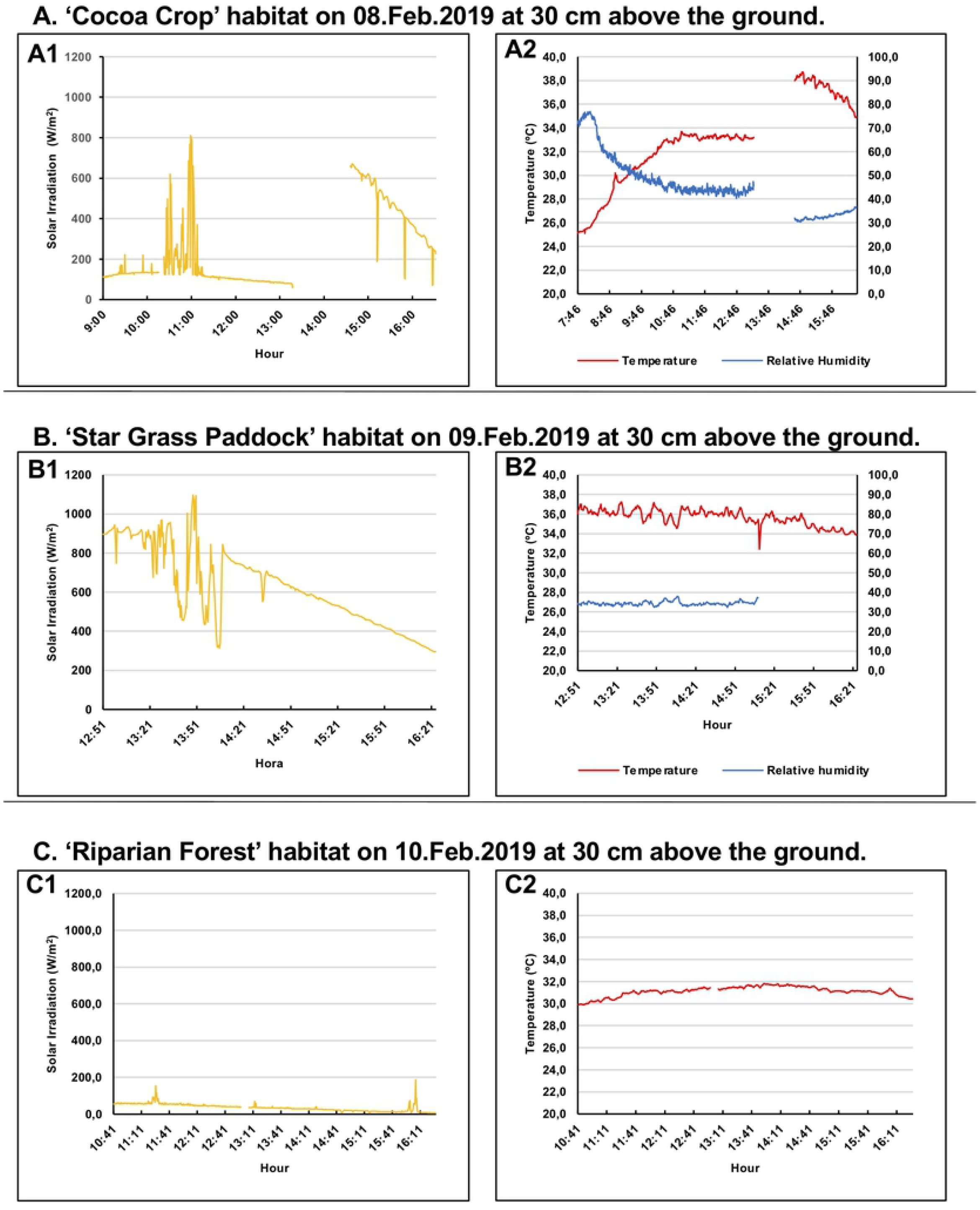
Plots of microclimatic variables that were measured *in situ* for three habitats. (A). ‘Cocoa Crop’ habitat. (A1). Solar irradiation values (W/m^2^) (N=736; 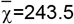; SD=189.1; Max=810.6; Min=59.4) according to the daytime hours. (A2). Temperature values (°C; red line) (N=901; 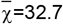; SD=3.6; Max=38.7; Min=25.1) and relative humidity (%; blue line) (N=901; 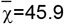; SD=11.6; Max=76.9; Min=30.4) according to the daytime hours. (B). ‘Star Grass Paddock’ habitat. (B1). Solar irradiation values (W/m^2^) (N=427; 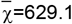; SD=199.6; Max=1,096.9; Min=295.6) according to the daytime hours. (B2). Temperature values (°C; red line) (N=427; 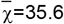; SD=0.8; Max=37.2; Min=32.4) and relative humidity (%; blue line) (N=276; 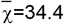; SD=1.0; Max=38; Min=32.3) according to the daytime hours. The relative humidity variable was not measured during the same period of time than temperature because of a damage in the respective sensor. (C). ‘Riparian Forest’ habitat. (C1). Solar irradiation values (W/m^2^) (N=677; 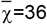; SD=19.8; Max=189.6; Min=6.9) according to the daytime hours. (C2). Temperature values (°C) (N=677; 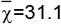; SD=0.4; Max=31.8; Min=29.9) according to the daytime hours. The relative humidity variable was not measured because of a damage in the respective sensor.

The relative humidity varied by 40% from the first part of the morning until midday, but in the afternoon, it increased by almost 10% from 30%. Thus, the relative humidity can vary up to 50% in a single day in the habitat of ‘Cocoa Crop’ (Figure 5). This one-day variation was half of the variation recorded in seven years for temperature and 75% of the variation recorded multiyearly for humidity (Table 7). Ticks in this environment should survive extremely low humidity conditions of up to 30% when there is more solar radiation. The relative humidity decreased as the temperature increased. Higher values of relative humidity, early in the morning, coincided with observations of increased tick activity and abundance in the dry ice traps at the time of capture.

### Star Grass Paddock

Variations were between 1.100 and 300 W/m^2^, which means a difference of 800 W/m^2^, decreasing towards the sunset. On the other hand, the temperature remained high on average at 35°C, decreasing little in the afternoon hours. Therefore, the range of variation was lower (5°C) in the afternoon than that observed in the morning for the habitat ‘Cocoa Crop’ at the same time (Figure 5). The relative humidity in ‘Star Grass Paddock’ was low in summer, 34% on average and at least 32%, but equally the variation between extreme values was low, only 6% (Figure 5).

### Riparian Forest

In this habitat, solar irradiation was very low (one order of magnitude lower), with an average of 36 W/m^2^ and a variation between extreme values of 180 W/m^2^ (Figure 5). This low penetration of sunlight, due to the high coverage of the canopy, maintained a relatively constant temperature with an average of 31°C, varying only 2°C. Unfortunately, due to damage to the humidity sensor, the respective values were not recorded that day.

### Microclimatic variables by habitat in winter

#### Cocoa Crop

Starting with a peak of 800 W/m^2^ and then reaching a second peak above 600 W/m^2^, the solar irradiation remained relatively constant below 200 W/m^2^ most of the time monitored. However, the average irradiance was lower, almost half, than in summer (126 vs. 243 W/m^2^; Figures 6 and 6). The variation in temperature was also smaller in winter (just 5°C) when compared to the summer season. Likewise, the variation in relative humidity was also lower in winter with about 30% between the extreme values when compared to 47% in summer (Figure 6).

**Figure 6.**
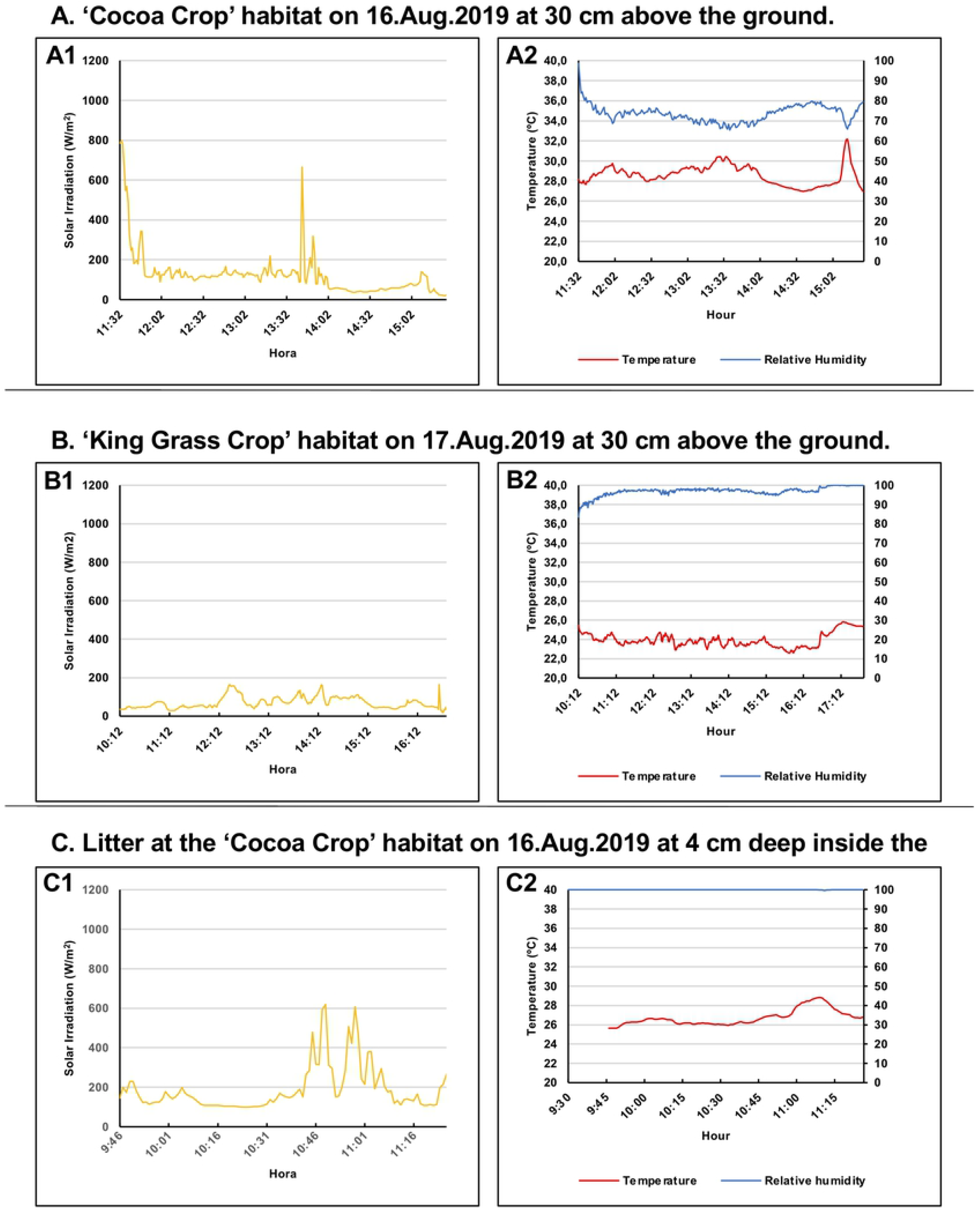
Plots of microclimatic variables that were measured *in situ* for two habitats. (A). ‘Cocoa Crop’ habitat. (A1). Solar irradiation values (W/m^2^) (N=236; 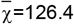; SD=118.3; Max=798.1; Min=19.4) according to the daytime hours. (A2). Temperature values (°C; red line) (N=236; 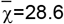; SD=1.0; Max=32.2; Min=27.0) and relative humidity (%; blue line) (N=236; 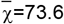; SD=4,3; Max=98.6; Min=65.4) according to the daytime hours. (B). ‘King Grass Crop’ habitat. (B1). Solar irradiation values (W/m^2^) (N=396; 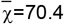; SD=30.2; Max=164.4; Min=19.4) according to the daytime hours. (B2). Temperature values (°C; red line) (N=457; 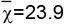; SD=0.7; Max=25,8; Min=22.6) and relative humidity (%; blue line) (N=457; 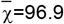; SD=2.3; Max=100; Min=83.6) according to the daytime hours. (C). Litter at the ‘Cocoa Crop’ habitat. (C1). Solar irradiation values (W/m^2^) (N=101; 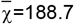; SD=114.1; Max=619.4; Min=99.4) according to the daytime hours. (C2). Temperature values (°C) (N=101; 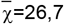; SD=0,8; Max=28,8; Min=25,6) and relative humidity (%) (N=117; 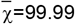; SD=0.05; Max=100; Min=99.6) according to the daytime hours.

### King Grass Crop

Solar radiation was low with an average of 70 W/m^2^ and a difference of 145 W/m^2^ between the extreme values recorded (cloudy day), which can be explained by the height of the King Grass (about 2 m) and the dense vegetation. In this sense, the humidity was very high at 30 cm from the ground, between 83-100% with an average of 97% during that day (Figure 6). The range of temperature variation throughout the day was also narrow with only 3°C and an average of 24°C.

### Litter in Cocoa Crop

The variables solar irradiation, temperature and humidity inside the litter, that is, between the soil and the litter, were taken in order to compare the litter (a place used as a refuge by ticks because its lower micro-environmental variation) with the ‘Cocoa Crop’ microenvironment at 30 cm from the ground. Thus, the average temperature inside the leaf litter (7 cm deep) was two degrees lower than in the environment, and the variation between extreme values was narrower. This suggests more stable and less extreme environmental conditions for the ticks that inhabit the litter. For its part, the humidity was maximum within the litter, almost completely constant (100% for most of the period studied), and 26% lower than the average of the microenvironment (Figure 6). Although the measurements were not simultaneous between the litter and the microclimate at 30 cm from the ground, the solar irradiation remained below 200 W/m^2^ most of the time in both environments, indicating that sunlight reaches the leaf litter at a depth of 7 cm.

Integrating the information from the sensors in all the habitats, with the aim of recreating the behavior of the variables in a day, it can be pointed out that the environmental temperature follows a normal distribution curve (except if it is cloudy or raining). The relative humidity shows a negative exponential behavior during the day (except if it rains, where the humidity exceeds 96%). The solar irradiation was inversely proportional to the vegetation cover, i.e. the greater the tree canopy and the greater the number of layers (Table 1), the less solar radiation will be registered by the sensor at 30 cm from the ground. Thus, the average solar radiation captured in summer in the habitat of ‘Star Grass Paddock’ was 2.5 times higher than that measured in ‘Cocoa Crop’ and more than 17 times higher than in ‘Riparian Forest’. In the same sense, the average room temperature was maximum in ‘Star Grass Paddock’ with 35.6°C, being 3°C lower in ‘Cocoa Crop’ and more than 4°C lower in ‘Riparian Forest’. Therefore, the climatic and microclimatic data show that, within the Matepantano farm, *A. mixtum* lives in an environment with large daily, seasonal and multiannual fluctuations in relative humidity, ambient temperature, solar radiation and precipitation.

### Potential Hosts

A total of 20 mammals were identified as potential hosts on the farm, which they have seen moving through the four habitats compared. 85% (17/20) of the animals were seen in ‘Riparian Forest’, 40% (8/20) in the area of administrative and residential buildings, 25% (5/20) in ‘Star Grass Paddock’, 20% (4/20) in ‘Cocoa Crop’, and 15% in ‘King Grass Crop’ (Figure 7). Only 3 of the 20 animals were classified as domestic and the rest were wild mammals. Armadillos, chigüiros, horses and cattle were identified by respondents as the species most frequently seen passing through the farm and connecting all of the four habitats. Six species of animals (three domestic and three wild) were the most seen mammals. Apart from the three species of domestic animals (cat, horse and cow), 14 species of wild mammals in summer and 12 in winter were consistently observed by respondents. It is interesting to note that summer was the time with the highest tick capture in our study.

**Figure 7.**
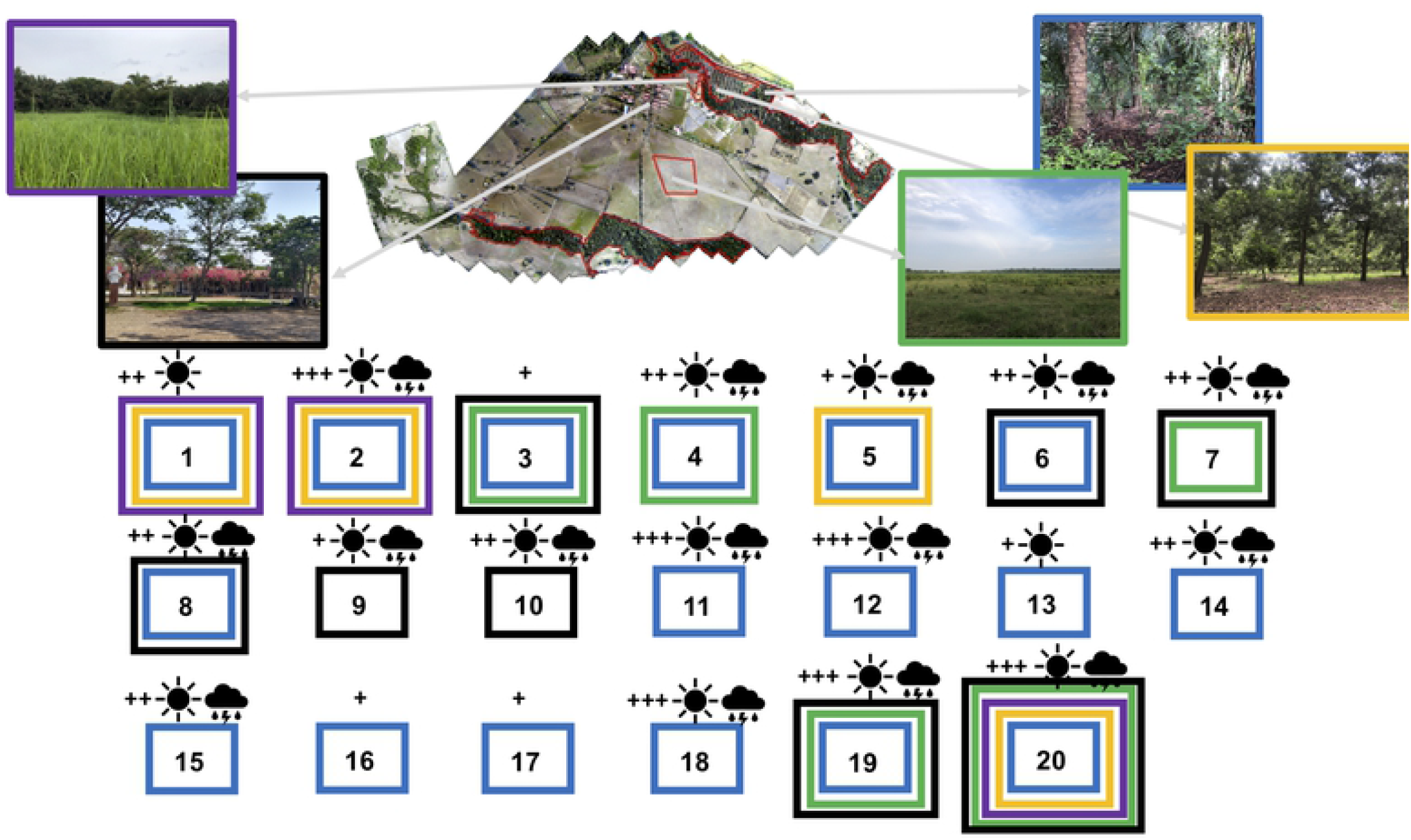
Potential host mammals of *A. mixtum* in the habitats of the Matepantano farm. Color of every box (blue = Riparian Forest; orange = Cocoa Crop; violet = King Grass Crop; green = Star Grass Paddock; black = administrative buildings; bedrooms, classroom, and restaurant) represent every habitat and other areas where the potential host has been seen by locals. Also, the sun symbol represents host presence in the summer season, while the rain clouds stands for host presence in the winter season. The ‘+’ symbol indicates the minimum perception of relative host abundance in the habitat, while the ‘+++’ represents the maximum perception. Note that host could have been seen in different habitats (several colored boxes). The animals species identified by locals based on photographs and common names have been identified by numbers and organized by rows from the left to the right and their common and scientific names (the last one between parentheses) are written as follows: 1) Armadillo (*Dasypus novemcinctus*); 2) capybara (*Hydrochaeris hydrochaeris*); 3) porcupine (*Coendou prehensilis*); 4) black agouti (*Dasyprocta fuliginosa*); 5) deer (*Odocoileus virginianus*); 6) flat-faced fruit-eating bat (*Artibeus planirostris*); 7) anteater (*Myrmecophaga tridactyla*); 8) Seba’s short-tailed bat (*Carollia perspicillata*); 9) fox (*Urocyon cinereoargenteus*); 10) opossum (*Didelphis marsupialis*); 11) tufted capuchin (*Cebus apella*); 12) Colombian red howler (*Alouatta seniculus*); 13) collared peccary (*Pecari tajacu*); 14) South American tapir (*Tapirus terrestris*); 15) southern tamandua (*Tamandua tetradactyla*); 16) puma (*Puma yagouaroundi*); 17) margay (*Leopardus wiedii*); 18) cat (*Felis catus*); 19) cow (*Bos taurus indicus*); 20) horse (*Equus ferus caballus*). Except for the last three species, all 17 animals seen by locals in the habitats are wild species. Both domestic and wild animals use to be around the administrative buildings.

## Discussion

The literature regarding empirical data on tolerance of *A. mixum* to different abiotic variables is still incipient. Even so, the available data about *A. mixtum* in Texas (USA), Mexico and Panama indicate that the species, in the field as well as in the laboratory, can tolerate high and low relative humidity, as well as high and low temperature (ranges: 35-95% and 18-36°C; see S5 Table). The thermal tolerance reported for *A. mixtum* coincides with our observations where the temperature fluctuates strongly between the two times of the year studied (Dry season (February/2019) vs. rainy season (August/2019)). In the Yopal region, the average temperature has been between 21 and 33°C for three decades (Table 6) and specifically at the Matepantano farm, between 18 and 37°C, for the last seven years (Table 7). We found that *A. mixtum* achieved the maximum niche width in summer, with occupation of all selected habitats at microclimate temperatures (in one day) ranging from 25 to 38°C (Figure 5). In this way, the population of *A. mixtum* that we study could be favored by the wide temperature changes that occur at different time scales (e.g. daily, monthly, seasonal and multi-year) in the habitats.

In the same way, one model projected that *A. mixtum* in California does not find any suitable habitat for the establishment of its populations, due to two related limitations, the low average temperature of 10°C associated to high altitude [42]. Additionally, [80] did not find *A. mixtum* above 1,200 m.a.s.l. in Panama, in places with less than 15°C. The thermal limit has also been corroborated by [81] where 50% of the adults of *A. cajennense* s.l. (renowned *A. mixtum*) died after 180 minutes of exposure to −12.5°C, limiting for this group their spatial distribution in latitudes higher than 30° [82]. From the above, it is inferred that the appropriate temperature in the habitats projected, used or preferred by *A. mixtum*, can vary widely within extreme values (> 10-15 °C) where the species is favored. The elevation of the terrain is undoubtedly associated with the thermal gradient that in turn determines the habitat that the species can occupy. The altitudinal distribution of *A. mixtum* in the foothills of the Eastern Colombian Mountains and its negative association with low temperatures, despite the presence of favorable hosts such as horses and cows, should be investigated to understand the distribution of the species in Colombia and other countries.

On the other hand, the adaptation or phenotypic plasticity of *A. mixtum* to relative humidity and temperature have important effects on their spatial distribution and habitat selection. Because of its wide tolerance to these two variables, the geographical distribution of *A. mixtum* extends from South Texas to the Pacific coast of Ecuador [48]. Additionally, [43] modeled the spatial distribution of *A. mixtum* from the records of presence-absence in the Neotropics and projected the presence of this tick in areas with high temperature variability (including the Colombian Llanos Orientales), preferring sustained high temperatures, within a narrow range of average annual temperatures (~19-33°C). The Estrada-Peña *et al*. model [43] projects the habitat of the semi-desert madrense of Mexico as not suitable for the establishment of *A. mixtum*. According to LandScope America (Available at: http://www.landscope.org/explore/natural_geographies/divisions/madrean_semidesert/), the Madrense semi-desert, which covers southern Texas and a substantial part of northern Mexico, has an average annual temperature of ~23°C and an annual precipitation of only 112 mm in the town of Brownsville (Texas).

The other key variable for considering a suitable habitat for *A. mixtum* has to do with the relative humidity (RH). [81] showed that no female *A. cajennense* s.l. (renowned *A. mixtum*) survived more than 1 month in very dry environments (35% of RH). Large fluctuations in relative humidity are reported in our study region. The extreme minimum value recorded in 29 years for the Yopal region was 61% and up to 30% relative humidity at Finca Matepantano (Tables 6 and 7, respectively). Values below 40% relative humidity are not usually common or prolonged over time and there are no data on their effect on the development, survival and population size of *A. mixtum*. [41] observed that prolonged exposure to 56% relative humidity stimulated quiescence in some stages of this tick. [81] recorded the death of all adults at 35% relative humidity and 33°C at the end of 30 days (S5 Table). In our study in Matepantano farm, we found that the relative humidity fluctuates strongly within a few hours in the same day and between seasons.

In summer, in the ‘Cocoa Crop’ habitat the relative humidity was low with an average of 46% and very low in ‘Star Grass Paddock’ with an average of 34% (Figure 5). In contrast in winter, the relative humidity was high for ‘Cocoa Crop’ with 74% average and very high for ‘King Grass Crop’ with 97% average (Figure 6). This variable could be related to population density and stage relative dominance. So, for example, we capture many more immature than adults from *A. mixtum* in summer, but no immature and almost twice as many adults in winter in ‘Cocoa Crop’ (Figure 3). However, in ‘Star Grass Paddock’ the capture of immatures was about 20 times higher than that of adults in summer; on the contrary, about five to 10 times less immature than adults were captured in ‘King Grass Crop’ in winter (Figure 3).

The results lead us to infer that a particular habitat is heterogeneous both in time and space and that the leaf litter on the ground could help to minimize the low humidity of the environment, since inside the leaf litter we measure higher relative humidity with respect to the environment. It also makes us think that larval and nymph stages could be more tolerant in summer at low relative humidity and high temperature, corroborating the great niche amplitude. However, this hypothesis still needs to be validated as well as to verify the response of adults in winter.

The contrasting readings of solar irradiation between ‘Star Grass Paddock’ and ‘Riparian Forest’ and its effect on the recorded values of relative humidity and temperature, as well as the lower capture of immatures in the first habitat in the summer season (Figures 3 and 5), coincide with the observations recorded in the literature. Higher relative humidity at ground level would be associated with increased foliage density and decreased sunlight penetration, which could be related to the preference of *A. mixtum* by areas of dense vegetation compared to open areas with short grasses [83,84]. In hydrophilic species of ixodids, such as *D. variabilis* o *I. uriae*, sub-adult stages (larvae and nymphs) tend to be more prone to water stress from drying out than adults [85,86]. However, *A. mixtum* is considered one of the most dehydration-tolerant ticks of ixodids [41], which would indicate its xerophilic adaptation.

Therefore, its tolerance to high temperatures and low relative humidity would explain its high population abundance in summer in the Matepantano farm. Where environmental conditions are desiccant or limiting, the microhabitat inside the litter (p. ej., 7 to 10 cm below the ground) could represent a refuge for ticks. This could explain why the low HR recorded in ‘Star Grass Paddock’, in summer, coincided with a higher capture of immature *A. mixtum*, when compared with winter and with respect to the total number of adults. The mechanisms to explain how each of the stages of *A. mixtum* conserving body water should be investigated.

According to [31,87], leaf litter is often a shelter for ticks in extreme weather conditions, particularly because its structure retains moisture and provides a stable temperature for ticks. In addition, potential ground hosts for tick sub-adults [88,89] may find food or protection in the leaf litter. The leaf litter of ‘Cocoa Crop’, presented an extreme humidity in winter, which could explain the absence of immature *A. mixtum* in that habitat during collection (Figure 6). On the other hand, in ‘Riparian Forest’ we observe a poor layer of leaf litter in summer and winter, probably due to the dominant vegetation (palms) or a high rate of decomposition. This, together with the water-saturated soil in winter could explain, in part, the low density of immature *A. mixtum* in this habitat. According to [90], the transformation, fragmentation of dense canopy forest (p. ej., deciduous trees) to those vegetal ecosystems with scarce canopy and less layers, it favors the flooding in winter.

In contrast, in summer, the low structural complexity of grassland vegetation, observed at the Matepantano farm, facilitates direct solar radiation, which rapidly dries out the top layer of soil, deteriorating the microclimate, increasing air temperature and decreasing relative humidity. Therefore, the more developed the vegetation in the landscape (strata, diversity, coverage, height) the less variation in environmental conditions and the lower the probability of extreme events of temperature, humidity and light radiation, which would favor the tick population dynamics. We hypothesize that the immature stages of *A. mixtum*, in the selected habitats of the Matepantano farm, are less susceptible to drying out when compared to adults in the summer season.

This is consistent with the findings of [91], where the nymphs of *A. sculptum* in Pantanal (Brazil) were found in the dry season, but not in the wet season. According to these authors, the absence of larvae and nymphs in the wet season in Pantanal can be explained by a seasonal pattern, which is analogous to the rainy season tick abundance in our study. [26,52] have observed the synchronization of the development (behavioral diapause) of a stage of the *A. cajennense* s.l, so that the next stage will take place under favorable weather conditions according to the season. The immature-adult ratios we observed were relatively similar, depending on the time of year, to those presented by [26,52].

Although we do not evaluate the population abundance of the *A. mixtum* free-living stages in all months of the year at the Matepantano farm, our observations present marked contrasts in the abundance of immatures between the summer and winter seasons in the selected habitats. The lower abundance or absence of immature stages of *A. mixtum* in the sampling carried out in the habitats ‘Cocoa Crop’ and ‘Riparian Forest’ in winter could be explained by: a) the higher soil moisture, which could promote a higher number of natural tick enemies [92–94] and b) the duration of the flooded ground (days). [95] point out that the seasonal flooding of soils in some regions of Brazil could be unfavorable to *A. sculptum*, tick that prefers drained soils in open areas with scrub and riparian forests.

If this occurs in the Matepantano farm, the presence of the three post-embryonic stages of *A. mixtum* suggests two possibilities: 1) there is only one generation per year and that they are successive, with a great reproductive peak between the end of the rains and the dry season. However, this hypothesis has yet to be validated; 2) there are overlapping generations (several per year), even if there is a reproductive peak generated by few females surviving the winter; also, because throughout the year there could be immature stages (as observed in the two contrasting periods) indicating several cohorts. This would be possible if the diapause capability of *A. mixtum* is activated at times of the year when their survival is threatened. In laboratory [96] found that engorged females from *A. auricularium* are resistant to water stress, when placed under immersion in distilled water for up to 96 hours, but the reduced progeny and the delayed oviposition time were attributed to diapause.

This would also help to explain how some females manage to survive harsh winters and flooding to maintain population size over time at the Matepantano farm. We observed that a few larvae of *A. mixtum* can survive up to one week under complete immersion in distilled water under laboratory conditions (unpublished information). It has also been shown that the eggs of *A. americanum* withstand underwater immersion periods of up to one week, without altering incubation time or hatching success [97].

For all the above reasons, we hypothesize that the peak of relative abundance of the immature stages of *A. mixtum* observed in summer at the Matepantano farm and, mainly, its depression in winter (Figure 3), could be the result in winter of: a) the direct death of some vulnerable immature stages, or b) the quiescent or diapause response of immature stages when experiencing extreme conditions of prolonged flooding and high relative humidity. In the first option, the adults survive the winter and lay eggs at the end of the rains, so some months later a high density of larvae and nymphs will be observed in summer. In the second option, the larvae and/or nymphs would go into diapause during the winter and come out the following summer.

In the first case there would be overlap of generations with a main reproductive peak. In the second case overlap of generations do not occur because of diapause. No doubt these hypotheses along with the ability of *A. mixtum* to survive long periods of time in flooded areas, particularly in years with La Niña phenomenon and high relative humidity, should be investigated, as well as their limits of tolerance to environmental variables related to extreme events. It is likely that each stage of *A. mixtum* tolerates and responds differently to abiotic variables and biotic interactions present in each habitat of the Matepantano farm. We also believe that individuals from *A. mixtum* would have the ability to select habitat, outside the host, as is the case with other tick species [85,98].

The wide tolerance of *A. mixtum* to abiotic variables is reflected in the tick population’s preference for multiple changing habitats, as we observed in our research. Therefore, we recommend investigating the mechanisms of habitat selection by this species, as well as the factors involved in the detachment of the engorged tick from the host when it perceives a habitat that is favorable to it, a process that we consider not to be random.

For its part, the fundamental niche of *A. mixtum* is determined by the tick’s tolerance range for the conditions it faces (i.e., temperature, humidity, solar irradiance), variables that define the breadth of habitat it colonizes. Thus, the niche width of *A. mixtum* is the range of habitats, with their range of resources, in which the growth rate of their population is positive. This niche amplitude increases from the genetic variation of the individuals with respect to the tolerance of the variables in the different types of habitat (e. g., differences in abundance between populations of stages observed in the Matepantano farm), and where generalist species increase their *fitness* as an adaptive strategy in unpredictable environments [99], as could be happening at the Matepantano farm or when the species faces interspecific competition [11].

Briefly, a generalist species is actually composed of different specialist individuals, called individual specialization [99]. This is an aspect we can explore further in different *A. mixtum* populations, since larvae had the highest niche width in summer, while adults had it on winter in our study. To understand niche width, it is necessary to investigate habitats with optimal niche, where *A. mixtum* reaches the maximum rate of population growth.

It makes sense for the tick species to have great niche amplitude since the longest period of its life cycle is outside the host [100] and can occur in different types of habitats. Conversely, a tick species specialized in one habitat type may be disadvantaged by habitat loss, fragmentation or transformation [101]. According to the above, the fragmentation of habitats in an agroecosystem like the Matepantano farm impacts the availability and diversity of habitats and hosts, and therefore the abundance of ticks at different times of the year, where the areas, cover and types of vegetation use to change. This fragmentation of the landscape [101,102] could be beneficial in the case of *A. mixtum* by offering a wide variety of environmental possibilities that favor their survival, reproduction and niche breadth.

Additionally, a large supply of hosts in the Matepantano farm and biological corridors, connecting patches of vegetation, would increase the connectivity between different habitats and expand the capacity of the species to be generalist by tolerating different conditions and having a wide range of resources to maintain the population and persist over time. The niche width of *A. mixtum* in the Matepantano farm was elevated at both times of the year for adults, which indicates its ability to resist environmental changes and to use different resources in changing or altered habitats, favoring its spatial dispersion by the wide range of hosts it uses [40,48,80,103]. However, the ecological niche of *A. mixtum* in the pre-adult stages (Figure 4) in the Matepantano farm was lower in the winter versus summer season, which may suggest less tolerance for variables such as flooding or less opportunity and availability to use the resources provided by the habitats.

This result certainly warrants further research. In summary, the three post-embryonic stages of *A. mixtum* have the capacity to use many resources and tolerate a wide range of environmental conditions in a highly heterogeneous agro-ecosystem in the Matepantano farm. Thus, *A*. *mixtum* could present adaptations, phenotypic plasticity and behavior modification [11] according to the habitat used, particularly those unstable (e. g., disturbed by humans and frequently changing, as King Grass Crop and Cocoa Crop) which they use immediately in an opportunistic way, as they are easily dispersed by a wide range of hosts.

The preference of the larvae and nymphs of *A. mixtum* by the habitats ‘King Grass plantation’ and ‘Star Grass paddock’ in both seasons of the year may be related to their unstable nature due to anthropogenic activities such as cutting and grazing (Figure 4). For example, the habitat ‘King Grass plantation’, composed of non-perennial grass *Pennisetum hybridum*, is temporary and its use and preference by *A. mixtum* is opportunistic. Our observations coincide with the hypothesis of physiological plasticity indicated for the *A. cajennense* s.l by [100], where the survival of such ticks can be explained by a set of habitat variables, and not by strict host specificity.

In contrast, the ‘Riparian Forest’ habitat was avoided by ticks according to the calculated use and preference rates (Figure 4 and Table 4). The explanation for this result could be related to a greater number of natural enemies; the constant export of ticks by wild animals to neighboring habitats; the relative climatic stability of the habitat; some factor that exceeds the tolerance of the ticks; or a system that has many natural enemies or controllers. Such hypotheses, however, must be valid.

In the same way, we notice that the abundance (population size *proxi* variable) went from high values in summer to a drastic decrease in winter (one order of magnitude), related to the season of heavy rains and floods. The question remains as how to explain the persistence of *A. mixtum* in the Matepantano farm, as the winter phenomenon could drastically affect the population size in the following year. To solve this question, we calculated the intrinsic rate of population growth = R (n_t+1_/n_t_), that relates the population abundance (n) in a time (t+1) with respect to a previous time (t); being R=1 a stable population, >1 growing and <1 in decline. Thus, if we divide the total abundance values quantified in winter (266) by those quantified in summer (6,733) for *A. mixtum* would give us a R = 0.039, suggesting population decrease between summer and winter, for the same year. That is, a population loss rate of 96.1% (1 - 0.039; or 100% - 3.9%).

However, we quantified in winter 77 females, of which not all of them will manage to reach engorgement during the winter, and many eggs by probability will not survive or develop into larvae. Furthermore, assuming that the effective amount of larvae in the following summer is equal to the already quantified summer, we would have 2,533 larvae. This means that when calculating R between the winter and the expected summer of the following year it would give: 2,533 larvae next summer/77 females in winter = 32.89 = R (1-32.89 = 31.89), giving a population growth of 3,189% (31.89*100%). To know the net balance in the population growth between the summer of the quantified year and the summer of the following year, we multiply the values of R, so R 0.039 × 32.89 = 1.28 (R= 1 - 1.28 = 0.28 or 0.28 x 100% = 28%), resulting in a net population gain of 28%.

This calculation suggests that, even with massive loss of individuals in the winter season (one order of magnitude), the population would be increased by 28% the following year, explaining the testimonies by the inhabitants of the Matepantano farm about the persistence of the species over time (decades). This population persistance could be explained because a few engorged females are enough to produce thousands of eggs and larvae to keep a positive population size. Nevertheless, other possible explanation for the population to persist could be a continuous supply of individuals (rescue effect) from neighbouring habitats or a source population (with positive growth), so these new immigrants could be carried by host, being part of so-called source-sink population dynamics [11]. This hypothesis should be further explored.

From the evidence collected we consider that the functional connectivity of the fragmented agroecosystem landscape at the Matepantano farm, particularly between habitats of King Grass Crop’ with ‘Cocoa Crop’ and ‘Riparian Forest’ is made possible by the presence of small mammals (marsupials, armadillos and rodents) and capybaras (*Hydrochoerus hydrochaeris*) at the Matepantano farm (Figure 6). Thus, the dispersion of the tick in space and the connection between habitats will depend on factors such as the availability of hosts; the activity and home range of the host [46,104]; the time of transport on the host [105]; and the host’s immune defense mechanisms.

According to [40,48], the adult collection of *A. mixtum* in the Neotropics and Nearctics indicates that adults of *A. mixtum* can feed on various domestic mammals and wild species, and parasitize humans [80,103]. The connectivity between the studied habitats by the constant transit of diverse hosts could help to colonize temporarily new or hostile, transformed or marginal habitats, as well as to serve as a rescue mechanism for tick populations in unsuitable habitats, where the mortality rate exceeds the birth rate [106–108]. Thus, some hosts serve as a bridge between habitats, either because in such habitats those hosts find food [32,109] or refuge from potential predators [31]. In turn, ticks take advantage of this dispersal mechanism even when the primary hosts are absent [36] or have been anthropogenically restricted [110], seeing a possibility of expansion, as they are opportunistic on modified fragmented and crop-based systems [111].

In our study area, the connectivity between adjacent habitats (King Grass Crop, Cocoa Crop, Riparian Forest, and Star Grass Paddock) could also be given by the independent movement from the host of ixodid ticks, regarding such a movement is usually limited, depending on the species, habitat conditions and the need to search for a host. For example, [101] observed that non-fed larvae of *I. ricinus* have a poor dispersal capacity due to their gregarious behavior, while the range of movement of free living nymphs and adults of *A. americanum*, a tick that hunts its host, was between 17 and 23 m, where most of the adults moved less than 13 m [104].

For their part, the nymphs of *A. maculatum* disperse slowly (1.27 cm/day) and seek out their host by moving very close to the ground (3.6 cm high) [112]. Although there is no data on the horizontal movement of each *A. mixtum* stage, field experiments in Texas [41] indicated that in the vertical displacement, in search of hosts, a third of the nymphs exceeded 30 cm from the ground and 3% of the adults were found between 10 and 60 cm, with a maximum of 115 cm, suggesting their hunting strategy, to favor their dispersion and feeding. Less vertical movement has been reported in the related species *A. sculptum* in Pantanal (Brazil) [91] with 15 to 50 cm of vertical climbing. We need to investigate the abilities of *A. mixtum* in its different stages to move between habitats.

Our findings validate modeling results for *A. mixtum* of [43] in the Neotropics and [113] in Colombia, who point out different types of vegetation cover, as well as hot and dry seasons, as favorable for the presence of this tick. Because of its high thermal tolerance and niche amplitude, *A. mixtum* colonizes several types of habitats in the Matepantano farm that provide diverse resources and climatic conditions, which coupled with functional connectivity by the hosts, allow it to adapt and survive in changing habitats. However, this characteristic of the tick represents a serious problem for the human and veterinary public health systems.

We consider, based on the results of our study, that *A. mixtum* is more vulnerable in winter, especially in the flooding season. In this regard, control measures should focus on the elimination of *A. mixtum* female in winter and the control of larvae and nymphs in habitats changed by human activity, with temporary or unstable characteristics around ‘Riparian Forest’, such as ‘Star Grass Paddock’ and ‘King Grass Crop’. Reducing the risk of rickettsial infection in humans should be the goal of control measures on the Matepantano farm and in the Llanos Orientales region.

It is recommended to monitor the populations of *A. mixtum* free-living stages in the different habitats of the Matepantano farm, as well as their parasitic populations on domestic and wild animals, in relation to the climate and microclimate at different times of the year. Such information will be useful in determining the local distribution of *A. mixtum* and to identify those key factors that explain the tolerance of the tick to extreme values of temperature and RH, in order to plan effective and specific strategies of control of its off-host stages in the region.

## Conclusion

*A. mixtum* presented preference for diverse environments in the Matepantano farm, as well as niche amplitude and tolerance to strong environmental fluctuations. All stages of *A. mixtum* use the ‘Riparian Forest’ but do not prefer it, which is compensated by the preference of three changing and transformed habitats in their surroundings. The population was more abundant in the dry season (larvae and nymphs) and decreased by one order of magnitude in winter (more adults, relatively). In winter, the immature stages were less abundant and could be more susceptible to the upper end of humidity and flooding, so control could be more effective there.

Further research should be developed to understand the morphophysiological or behavioral strategies and adaptations of *A. mixtum* to tolerate wide fluctuations and extreme changes in climate and microclimate. Several variables that falling within the scope of ‘functional habitat’ (e. g., host richness within each habitat associated to *A. mixtum;* host parasitic load; host biotic interactions with natural enemies; and host role in landscape connectivity, maintenance and persistence of tick populations) should be included in future research. Also, tick intrinsic variables, like photoperiod behavior and development, might help to explain the parasitic and non-parasitic population peaks of all *A. mixtum* stages in dry and wet seasons, to determine generational overlap, and to distinguish strategies for survival for every developmental stage. The population control of *A. mixtum* should be carried out at the time of greatest vulnerability (e. g., winter), where the population size is low, and must be directed mainly towards females.

## Acknowledgements

We thank Sergio Andrés Viniescas Pineda and Victor Montaña, responsible for the Davis Weather Station, (Finca Matepantano, Yopal, Casanare) for sharing with us the weather data of seven consecutive years. Also, we are grateful for the immense support of Patrick Nicolás Skillings for taking and processing the images of the Matepantano farm with drone. Finally, our recognition to the undergraduate students in Veterinary Medicine of La Salle University Laura Coy and Rafael Salazar, and to the graduate students in Biology of the Pontificia Universidad Javeriana Náyade Cortés and María Fernanda Cogollos. Our special thanks to the Young Researcher Juliana Gil, sponsored by Colciencias.

## Author contribution statement

A.A., E.B. and M.H. conceptualized the project. M.H. received research funds from the Pontificia Universidad Javeriana, Bogotá Campus. A.A., E.B. and E.F.-B. carried out the field work. A.A. conducted, analyzed and presented the statistical information of habitat use and preference in tables and figures, and structured the document with the support of E.F.-B. and E.B. A.A. and E.F.-B. wrote the manuscript. All the authors reviewed and edited the manuscript in its different sections.

## Contribution to the field statement

Empirical data of high niche amplitude in an agroecosystem with fragmented habitats and availability of a variety of domestic and wild hosts represent new knowledge for the species.

## Supporting information captions

**S1 Appendix. Procedure used for drone orthomosaic mapping.**

**S2 Appendix. Procedure for vegetation cover estimation within each habitat.**

**S3 Appendix. Description of the making process for the ice traps and for the white flannelettes to collect free-living ticks on the field.**

**S1 Table. Codification for every ice trap (N = 24) with effective tick collection in summer, including sample code, habitat and GPS coordinates.**

**S2 Table. Codification for every transect (N = 22) with effective tick collection in summer, including sample code, habitat and GPS coordinates.**

**S3 Table. Codification for every ice trap (N = 31) with effective tick collection in winter, including sample code, habitat and GPS coordinates.**

**S4 Table. Codification for every transect (N = 22) with effective tick collection in winter, including sample code, habitat and GPS coordinates.**

**Table 8. Abiotic variables recorded for *Amblyomma mixtum* in literature.**

